# Organization principles of dynamic three-dimensional genome architecture associated with centromere clustering states

**DOI:** 10.1101/2025.06.12.659004

**Authors:** Satya Dev Polisetty, Shuvadip Dutta, Rakesh Netha Vadnala, Dimple Notani, Ranjith Padinhateeri, Kaustuv Sanyal

## Abstract

Fungal centromeres are clustered near microtubule organizing centers to help adopt the Rabl chromosomal organization. The role of centromere clustering in driving large-scale changes in structural and functional chromatin assembly remains unclear. Here, using Hi-C and super-resolution microscopy, we show that cell cycle-dependent centromere declustering and clustering states in *Cryptococcus neoformans* drive global changes in the 3D genome architecture. Centromeres and telomeres are scattered around the nuclear periphery at interphase^G1^, and this arrangement constrains the inter-arm interactions within a chromosome, providing a unique interphase^G1^ chromosome organization. Moreover, centromeres and telomeres are organized as compartments, segregating them from active euchromatic regions. Polymer modeling reveals that the transition from the unclustered to clustered centromere state during the cell cycle involves changes from a globular to an elongated chromosome architecture. Strikingly, while clustered centromeres replicate early in most yeasts, *C. neoformans* centromeres replicate late in S-phase, hinting a possible link between centromere clustering dynamics and *CEN* DNA replication timing. Overall, our study uncovers several unique organizational principles governing the dynamic genome architecture in an evolutionarily diverged basidiomycete yeast.

**Significance:** Chromosomes occupy the nuclear space in many ways. Primary chromosomal arrangements are such that centromere regions of different chromosomes either form a cluster, as in yeasts, or are scattered around the nuclear periphery, more common in metazoans. Exceptionally, centromeres show cell cycle stage-specific clustering in the basidiomycete fungus *Cryptococcus neoformans*. We show that the spatial positioning and the refractory nature of centromeres and telomeres shape the arrangement and large-scale organization of chromosomes in *C. neoformans*. Metazoan-like late-replicating centromeres in *C. neoformans* possibly favor the unclustered/scattered centromere state in S-phase, not commonly found in fungi. Our results not only highlight the remarkable genome plasticity of *C. neoformans* but also raise the possibility that centromere replication timing determines genome organization principles.

## Introduction

Chromosomes in a eukaryotic cell are three-dimensionally organized in two recurring configurations based on the spatial positioning of centromeres and telomeres^1^, Rabl and non-Rabl/Type-II. In the Rabl configuration, centromeres and telomeres are spatially separated within the nucleus and arranged in separate clusters. The clustered centromeres and the telomere cluster are located at the nuclear periphery, but these two clusters are spatially positioned on opposite ends of the nucleus. In contrast, the non-Rabl/type-II configuration is marked by the territorial arrangement of chromosomes, where individual centromeres and telomeres are scattered across the nucleus, sometimes forming subclusters. Well-studied fungi in the phylum of Ascomycota show the Rabl arrangement of chromosomes^2^, but most metazoan chromosomes tend to adopt the non-Rabl configuration^2^. Strikingly, *Schizosaccharomyces pombe* and *Magnaporthe oryzae* genomes adopt the Rabl arrangement^3,4^ except transient declustering of centromeres is observed at metaphase. In the fruit fly *Drosophila melanogaster*, the centromere clustering is developmental stage-specific. While chromosomes in embryonic nuclei display the Rabl configuration^5^ with clustered centromeres, more differentiated fly cells, such as spermatocytes^6^ have unclustered centromeres as chromosomes adopt the non-Rabl organization. Metazoans, including humans, exhibit differentiation-specific clustering of centromeres with the nucleolus, which is lost in terminally differentiated cells^7^. Intriguingly, centromeres show a strong propensity to cluster around the nucleolus in human cancer cells^7^ although the role of centromere clustering in cancer cells needs to be explored.

Apart from shaping the chromosome-scale architecture, spatial positioning and clustering of centromeres impact the regulation of several structural aspects of the genome, including the telomere bouquet formation during meiosis in *S. pombe*^*8*^. While the importance of centromere clustering in shaping the genome architecture and function is significant, functional and architectural differences involved in clustered and unclustered centromere states are underappreciated. In this study, we report that in the basidiomycete budding yeast, *Cryptococcus neoformans*^9^, the centromere clustering states differ according to the cell cycle stages - centromeres are unclustered during interphase and gradually cluster during mitosis^9^. This unique feature of dynamic changes in chromosomal organization makes *C. neoformans* an attractive model to track the natural transition of two major chromosome arrangements that exist across species. While the spatial positioning of centromeres orchestrates the overall three-dimensional genome organization^10^, *C. neoformans* holds the key to link chromosome architectural changes associated with centromere clustering during the mitotic cell cycle.

## Results

### *C. neoformans* genome switches between non-Rabl and Rabl configurations

The cell cycle stage-dependent spatial clustering of centromeres/kinetochores has already been reported in *C. neoformans*^9,11^. We identified the homolog of Protector of Telomeres (Pot1), a single-strand telomere DNA-binding protein^12^ (Supplementary Fig. 1) in *C. neoformans*. GFP-Pot1 was expressed in a strain where centromeres were marked with mCherry-tagged CENP-A. Telomeres in G1 cells were visible as multiple puncta located at the nuclear periphery, similar to centromeres (Fig. 1a), unlike any polarized distribution observed in *Saccharomyces cerevisiae* or *S. pombe*^13-15^. Instead, centromeres and telomeres in *C. neoformans* show no prominent spatial clustering resembling interphase cells of metazoans such as humans^16^. Centromeres in *C. neoformans* appear to cluster during G2 and anaphase as chromosomes adopt the Rabl-like arrangement (Fig. 1a and b), typically observed in yeast species. While there are 14 chromosomes in *C. neoformans*, the number of telomere puncta observed in G1 cells is significantly less than 28, indicating telomeres form subclusters (Fig. 1c). We could not detect Pot1 signals in most metaphase cells when centromeres migrated into the daughter bud. Notably, while centromeres do not form a single punctum, 14 individual centromere puncta in each G1 cell could not be detected, indicating centromeres form subclusters^9^. To further resolve the spatial positions of centromeres and telomeres, we imaged cells using 3D Structured Illumination Microscopy (SIM^2^) (Fig. 1d). This approach revealed that centromeres and telomeres are indeed scattered across the nucleus and occupy non-overlapping spatially proximal positions in interphase^G1^ cells.

**Fig. 1:**
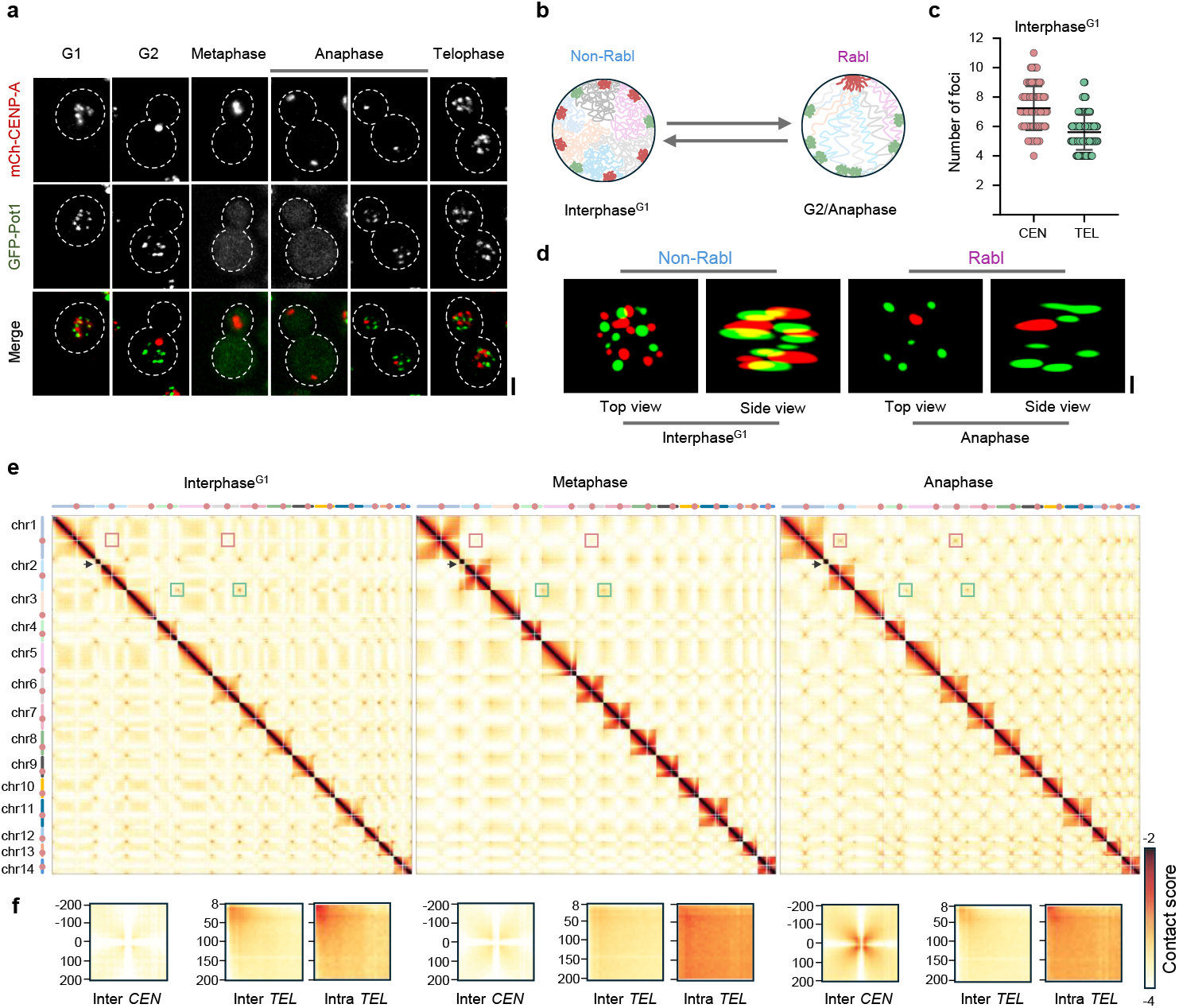
The *C. neoformans* genome alternates between the non-Rabl and Rabl configurations during the cell cycle. **a**, Microscopic images of centromeres and telomeres marked by mCherry-CENP-A and GFP-Pot1, respectively, at different cell cycle stages. Scale bar, 2 μm. **b**, Schematic of the non-Rabl and Rabl arrangement of chromosomes. **c**, Quantification of the number of centromere or telomere puncta in interphase^G1^ stage, n=90 cells. The mean values are represented as a black bar. Error bars represent SD. **d**, 3D rendering of microscopic images of centromeres and telomeres of indicated cell cycle stages, Scale bar, 0.5 μm. **e**, Whole genome contact maps of interphase^G1^, metaphase, and anaphase stages binned at 8 kb resolution. Chromosomes with centromeres marked in red circles are displayed on the left and at the top of the contact maps. Red and green squares on the maps represent inter-centromere and inter-telomere interactions, respectively. The black arrow marks the ribosomal DNA (rDNA) locus on chromosome 2. **f**, Aggregate Hi-C maps of inter-centromere/telomere and intra-telomere interactions (between left and right arms of the same chromosome), binned at 8 kb resolution, of all chromosomes. For calculating inter-/intra-telomere contacts, the left arm of the chromosomes was used. The color scale from yellow to brown represents low to high contact scores (log_10_), respectively. e and f panels share the same color scale.

### Interphase^G1^ genome is characterized by strong inter-telomere interactions

To study the cell cycle-associated changes in the genome organization of the *H99* strain of *C. neoformans*, we performed Hi-C in synchronized interphase^G1^ and mitotic cells at metaphase and anaphase (Supplementary Fig. 2 a, b and c), and generated whole genome contact maps (Fig. 1e). Hi-C was performed in two replicates at each of these stages. To assess the reproducibility of Hi-C data sets, we calculated pair-wise Pearson correlation coefficients of the datasets binned at 8 kb. The results indicate high concordance among the biological replicates (Supplementary Fig. 2d).

A striking feature of contact maps of a genome with the Rabl configuration of chromosomes is the presence of strong inter-centromere interactions^17-19^. Centromeres in *C. neoformans* are regional and span between 21 to 64 kb^20^. Inspection of Hi-C maps from synchronized interphase^G1^, metaphase, and anaphase cells of *C. neoformans* (Fig. 1e and f) reveals that features of the interphase^G1^ contact map are distinct from those of ascomycete yeasts possessing the constitutive Rabl organization^2^. Contact maps confirm that interphase^G1^ cells lack usual strong centromere interactions between the chromosomes, suggesting the non-Rabl metazoan-like chromosome arrangement (Fig. 1e, red squares and 1f and Supplementary Fig. 3a) in a fungal cell. In fact, inter-telomere interactions across the chromosomes are strongly enriched (Fig. 1e, green squares and 1f and Supplementary Fig. 3a). Such telomere-telomere interactions significantly reduce at metaphase, possibly due to chromatin compaction. The telomere interactions associated with left and right arms of the chromosomes remained uniform across stages (Fig. 1f, Supplementary Fig. 3a). While microscopic observations demonstrate the existence of multiple telomere puncta, Hi-C data support enriched telomere contacts, confirming subclustering of telomeres during interphase^G1^, and anaphase. Perhaps the subcluster composition of telomeres is heterogeneous at the single cell level, which explains prominent telomere-telomere interactions in contact maps since Hi-C contacts are population averages. The metaphase stage in *C. neoformans* is characterized by a modest increase in inter-centromeric interactions as compared to interphase^G1^, consistent with microscopy results where centromeres appear to be loosely clustered and stretched along the spindle during metaphase^9^. Hi-C data reveal that anaphase cells of *C. neoformans* exhibit features similar to the Rabl configuration, characterized by the strongest *trans*-chromosomal interactions mediated by centromeres as well as telomeres (Fig. 1e and f, Supplementary Fig. 3a).

### Nuclear positioning of centromeres and telomeres shapes global chromosome architecture

Having established that *C. neoformans* chromosomes could adopt both the non-Rabl and Rabl-like arrangements in a cell cycle stage-specific manner, we sought to explore changes in the chromosomal organization facilitating the transition. A comparison of percent *cis* versus *trans* contacts across cell cycle stages reveals that the majority of interactions in interphase^G1^ chromosome are *cis*, restricted within the same chromosome as less than 20% of the contacts across the chromosomes were between different chromosomes (*trans*) (Supplementary Fig. 3b). On the other hand, metaphase and anaphase chromosomes exhibit a modest increase in *trans* contacts as compared to interphase^G1^ (Supplementary Fig. 3b). To visualize the differences in chromatin contacts across stages, we plotted observed/expected maps (Fig. 2a). A comparison of contact maps across stages identifies that pericentromeric regions are refractory to interact with the rest of the chromosome (Fig. 2a, solid ovals), a feature conserved in budding and fission yeasts^18,19^. Notably, subtelomeric regions at interphase^G1^ also behaved similar to centromeres (Fig. 2a, dashed ovals). This indicates the refractory nature of both pericentromeric and subtelomeric regions constrain the inter-/intra-arm interactions, thus shaping the non-Rabl interphase^G1^ chromosome organization (Fig. 2a, below). Unlike interphase^G1^, subtelomeric regions at anaphase engage in extensive interactions with the arms of the same chromosome (Fig. 2a, dashed oval). Across the stages, interactions among the arms of a chromosome increased as the distance from the centromeres increased (Fig. 2a, black box). The arm-arm interactions originating from centromeres are more enriched at metaphase and anaphase, implying the folding of arms at centromeres (Fig. 2a, below). Such arm-arm interactions adjacent to centromeres are absent in the interphase^G1^ genome. Centromere-centered aggregate maps across stages yield a similar trend (Fig. 2b, black boxes). These differences further confirm that the folding of metaphase and anaphase chromosomes resembles the properties of chromosomes arranged in the Rabl form. These aggregate maps also indicate that the pericentromeric interactions within the arms are stronger in interphase^G1^ as compared to anaphase (Fig. 2b, dashed ovals), revealing these regions are highly compacted in interphase^G1^. In metaphase, the overall increase in interactions around centromeres may be due to compaction of chromatin in general. To test if the stage-dependent compaction changes are restricted only to pericentromeric regions, we analyzed aggregate profiles of telomeres belonging to the left arm and noticed that interactions along the diagonal remains similar across the stages (Supplementary Fig. 4). To confirm these observations, we plotted average compaction profiles for individual chromosomes across stages (Fig. 2c and Supplementary Fig. 5). Note that the compaction profiles highlight the extent of local compaction within a specific chromosome and thus a comparison across different Hi-C experiments may not be appropriate. We observed that the regions flanking centromeres are highly compacted as compared to the rest of the chromosome in interphase^G1^ and metaphase. Surprisingly, the extent of compaction of these regions at anaphase is on par with the chromosome compaction levels. In all three stages, namely interphase^G1^, metaphase, and anaphase, telomeres remained highly compacted.

**Fig. 2:**
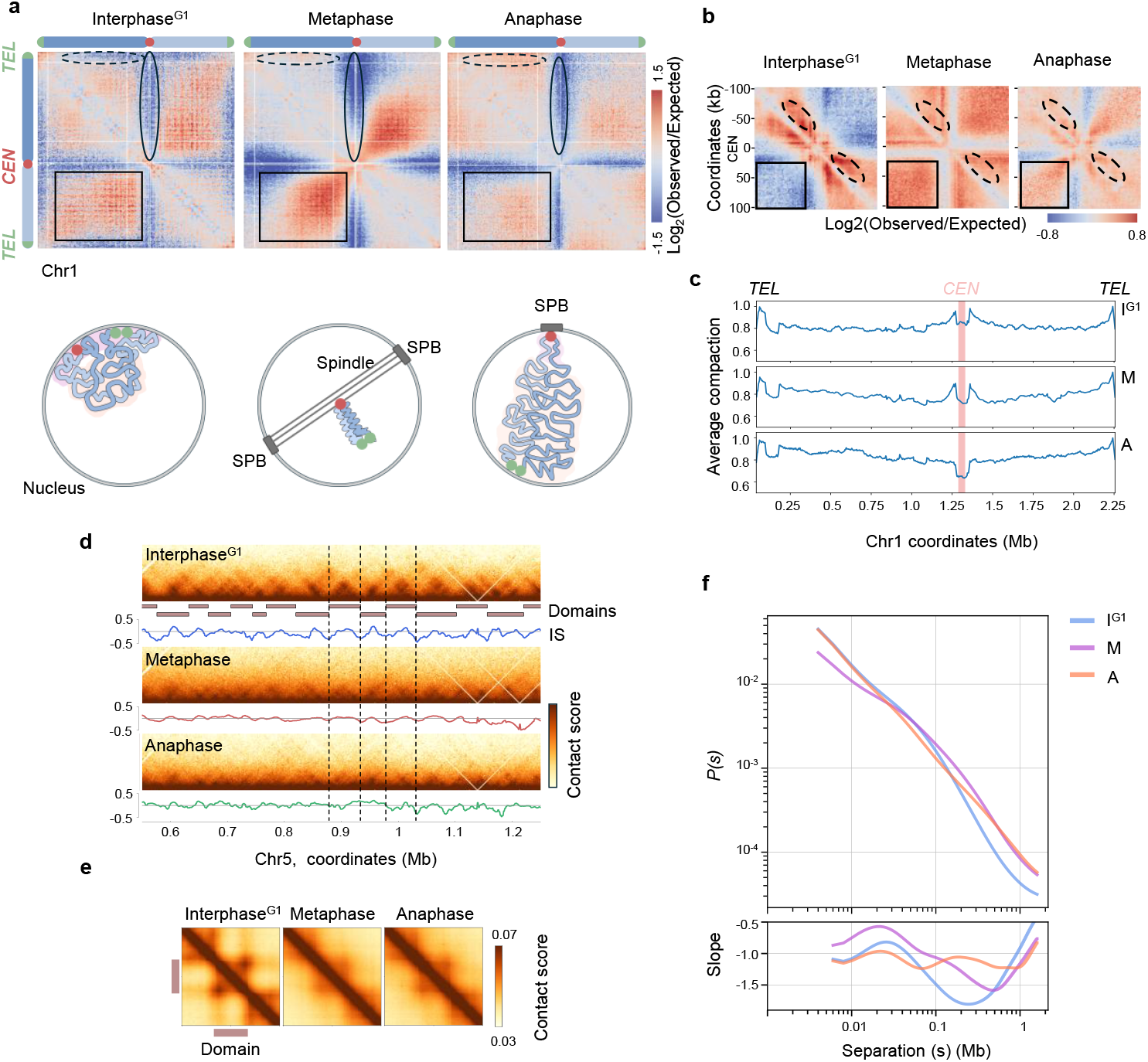
Transition from the non-Rabl to Rabl configuration involves global chromatin decompaction. **a**, *Top*, Comparison of observed/expected maps of chromosome 1 of interphase^G1^, metaphase, and anaphase stages, binned at 8 kb resolution. The blue to red color scale represents depletion and enrichment of contacts, respectively (log_2_ scale). Solid and dashed ovals on the maps denote the pericentromeric and subtelomeric regions, respectively. Black boxes highlight inter-arm interactions within a chromosome. *Below*, schematic highlighting structural changes in the chromosome organization across interphase^G1^, metaphase, and anaphase stages. **b**, Observed/expected aggregate Hi-C maps of *cis*-contacts around 100 kb regions, surrounding all 14 centromeres, binned at 2 kb. Color scale as indicated above. Dashed ovals represent contacts within the arms of the chromosomes around centromeres. Black boxes represent inter-arm interactions around centromeres. **c**, Compaction profiles of chromosome 1 for the indicated stages. The x-axis represents chromosome coordinates, and the y-axis represents average compaction. The red-shaded rectangle highlights the centromere position. **d**, A region of chromosome 5 Hi-C contact map (binned at 2 kb) of interphase^G1^, metaphase, and anaphase stages. Solid bars below interphase^G1^ contact map represent domains identified using interphase^G1^ Hi-C data. Insulation scores (IS) are indicated below each contact map. Dotted lines represent domain boundaries. The color scale from yellow to brown represents low to high contact scores, respectively (log_10_). **e**, Aggregate maps of domains across the three cell cycle stages (binned at 2 kb). The coordinates of the domain boundaries predicted using interphase^G1^ Hi-C data were used to generate the aggregate maps of metaphase and anaphase. **f**, *Top*, contact frequency *P(s)* curves plot of Hi-C data obtained from interphase^G1^, metaphase, and anaphase binned at 2 kb. *Below*, derivatives (d log P(s)/d log(s)) of the *P(s)* curves.

Next, we studied the differences in local chromatin packing across the cell cycle stages. Visual inspection of Hi-C maps at 2 kb resolution indicated the presence of strong self-interacting domains, similar to the domains reported in fission yeast^18^, in interphase^G1^ (Fig. 2d), To quantify changes in the domain organization, we determined domain boundaries by estimating genome-wide insulation scores^21^ from interphase^G1^ Hi-C data and generated aggregate Hi-C maps of identified domains (Fig. 2e). The analysis revealed the presence 40-70 kb long domain-like structures in interphase^G1^ chromosomes (Supplementary Fig. 6). The presence of these domains was significantly reduced during metaphase, likely due to the overall compaction of chromosomes. Such domains started to appear during anaphase, hinting at the gradual reestablishment of domain-like structures, a phenomenon observed in humans^22^. The striking resemblance of the disappearance of the domain-like structure across kingdoms of life is noteworthy. Based on these observations, we suggest that local as well as global chromatin architecture and the spatial positioning of centromeres and telomeres shape the cell cycle stage-specific chromosome reorganization.

To understand the difference in the nature of chromatin folding across mitotic stages, we plotted intra-arm contact probability (*P*) over the genomic distance, *s. P(s)* values change in a cell cycle stage-specific manner^23^. A power-law scaling between s^-1^ and s^-1.5^ indicates a fractal globule, and above s^-1.5^ indicates an equilibrium globule. The *P(s)* of interphase^G1^ uncovered two regimes of power-law scaling. Regions ranging from 10 to 100 kb exhibited ∼s^-0.93^ and those above 100 kb showed s^-1.72^ (Fig. 2f, below and Supplementary Fig. 7a). The observed power-law scaling during interphase^G1^ in *C. neoformans* indicates that at distances <100 kb, chromatin is organized as a fractal globule, and at >100 kb, chromatin exhibits the properties of an unconstrained polymer. The metaphase stage has a slower rate of decay at distances <100 kb with a power law scaling of s^-0.62^, indicating compaction of the chromosomes, and >100 kb, the decay was rapid with ∼s^-1.31^ (Fig. 2f, below and Supplementary Fig. 7a), which is similar to metazoan metaphase chromosomes^23^. Anaphase chromatin, rather strikingly, showed a uniform steep decay in contact frequency with values s^-1^ and s^-1.1^ for short and long distances, respectively (Fig. 2f, below and Supplementary Fig. 7a). This suggests that both at short and long distances, chromatin is folded as a fractal globule. The above-mentioned features were observed for the arms of individual chromosomes as well (Supplementary Fig. 7b-d). A comparison of interphase^G1^ and anaphase *P(s)* curves reveals that contacts between regions spanning 20-100 kb are enriched in interphase^G1^, and the interphase^G1^ domains are of sizes 40-70 kb. We speculate that the loss of contacts at 20-100 kb distances (Fig. 2f) perhaps corresponds to the observed loss of strong domain-like structures in anaphase (Fig. 2e). On the other hand, contacts spanning ∼ 150 kb are enriched in anaphase, providing an explanation for the observed increase in long-range contacts that encompass subtelomeric regions evident from O/E maps. To further confirm that anaphase chromosomes indeed exhibit enhanced distal/long-range contacts, we plotted the Distal Contact Index (DCI). DCI quantifies the ratio between distal and local contact strengths, providing a measure of the extent of long-range chromatin interactions. DCI analysis revealed an enrichment of distal contacts >70 kb across all chromosomes during anaphase as compared to interphase^G1^ (Supplementary Fig. 8a and b). The differences in the domain organization in interphase^G1^ and anaphase stages, combined with the changes in the decay pattern of *P(s)* curves, confirm that anaphase chromatin is less compact than interphase^G1^ with reduced short-range interactions but more prominent long-range interactions.

### Centromeres and telomeres form separate compartments in interphase^G1^

Since the pericentromeres and subtelomeres are refractory to interacting with the rest of the genome, we generated a correlation matrix^24^ to test if these regions behave similarly to the A and B compartments observed in metazoans^24-28^. Though we did not observe a plaid pattern that is present in metazoan contact maps^24^, we noticed that both pericentromeric and subtelomeric regions in *C. neoformans* remained in one compartment, referred to as the centromere-telomere (CT) compartment, and the rest of the chromosome remained in the non-CT compartment (Fig. 3a, left panel). This pattern was lost at anaphase, where only pericentromeric regions are segregated from the rest of the genome (Fig. 3a, right panel). Next, we performed eigendecomposition^24^ and the first eigen vector was used to estimate the positions of the compartments. The principal component (PC1) indicated the presence of two large compartment-like structures in all chromosomes, except for chromosomes 2 and 12. The CT compartments were enriched with H3K9me2 and H3K27me3 marks and have reduced transcription (Fig. 3b and c, Supplementary Fig. 9). To find out if these CT compartments physically interact among each other, we plotted interactions whose log_2_ observed/expected ratio is >3 using circos (Fig. 3b). Indeed, strong interactions were detected between subtelomeric regions of different chromosomes within the CT compartment but not with centromeres. These observations demonstrate that the *C. neoformans* interphase^G1^ genome is explicitly organized into two large euchromatic and heterochromatic compartments.

**Fig. 3:**
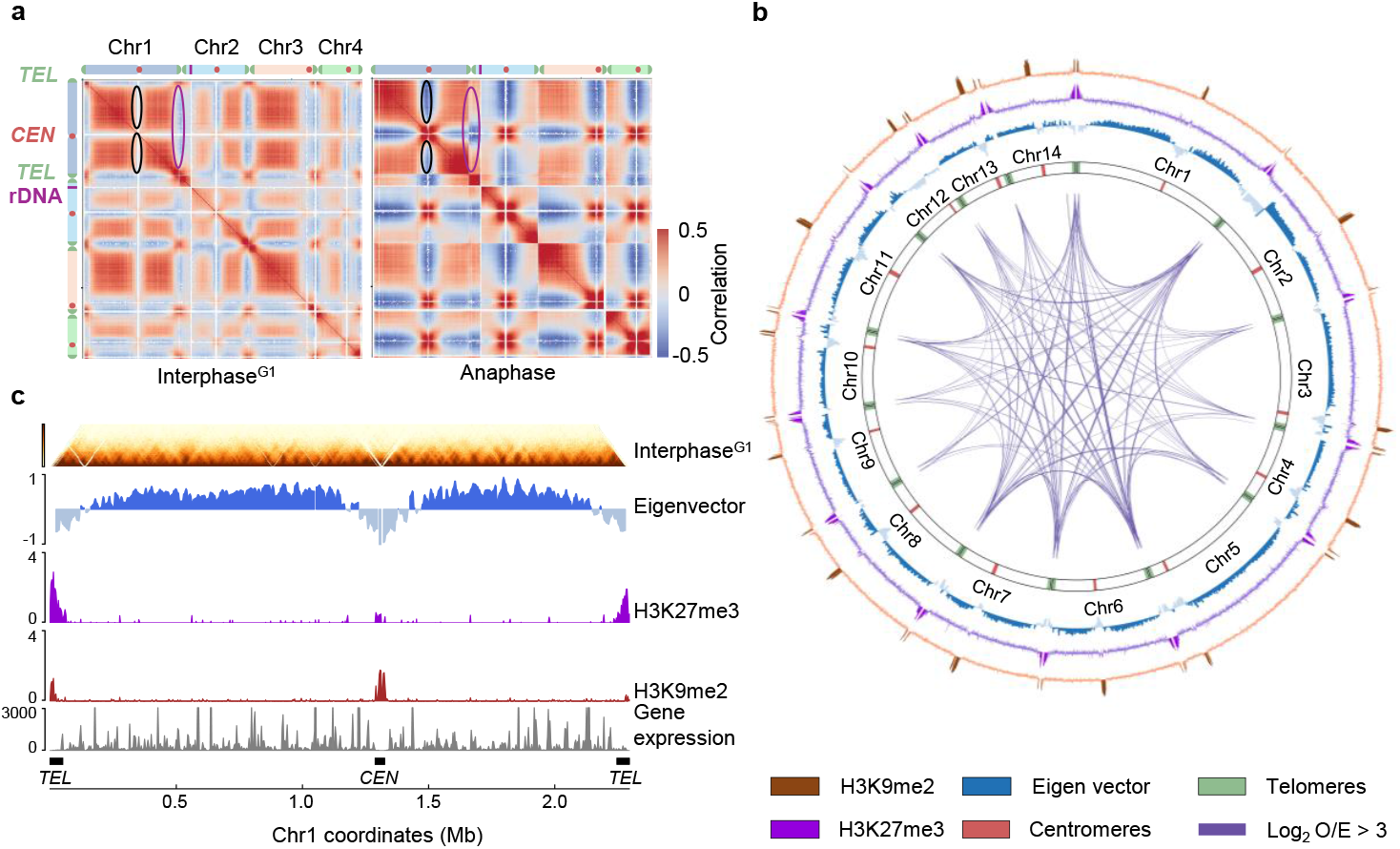
Two-compartment-like organization of the *C. neoformans* interphase^G1^ genome. **a**, Correlation matrix of contacts for chromosomes 1 – 4 in interphase^G1^ and anaphase stages, binned at 8 kb. The color map from blue to red represents negative to positive correlation values. Black and violet ovals represent pericentromeric and subtelomeric compartments. **b**, Circular (circos) plot representing the interactions across the subtelomeric regions and the associated chromatin modifications and eigenvector. Violet links represent the strongest inter-chromosomal contacts whose log_2_ (observed/expected) values are greater than 3. From outside to inside: H3K9me2 (brown), H3K27me3 (violet), eigenvector (blue), chromosomes marked with centromere (red) and telomeres (green) positions. **c**, Hi-C contact map of chromosome 1 binned at 2 kb at interphase^G1^, corresponding tracks representing eigenvector, log_2_ ChIP enrichment profiles of H3K27me3, H3K9me2, and gene expression levels are indicated below. Black bars represent centromeres and telomeres of chromosome 1. The color scale from yellow to brown represents low to high contact scores, respectively (log_10_).

### Modeling chromatin architectural states associated with clustered and unclustered centromeres

To better understand the overall 3D folding features of the chromosomes in Rabl and non-Rabl states, we developed a polymer model by coarse-graining the *Cryptococcus* genome into 14 flexible chains made of beads connected by springs, each representing one chromosome. Each bead represents a 10 kb chromatin segment, which is equivalent to about 50 nucleosomes. The polymer models were built in two sequential stages (Fig. 4a) to progressively reconstruct the global 3D genome organization. First, we generated an ensemble of individual chromosome conformations guided by Hi-C data. Next, chromosome-wise conformation states, the number of centromere and telomere clusters, and the nuclear volume were integrated to simulate the whole genome by placing all 14 chromosomes within a spherical confinement, mimicking the physical boundaries of the nucleus. Physical constraints due to stage-specific clustering of centromeres and telomeres (Fig. 4b) were introduced based on microscopy data (Fig. 1a). Visual inspection of the polymer models revealed that interphase^G1^ chromosomes form compact globular structures, whereas anaphase chromosomes are axially elongated and less compact (Fig. 4c and d). Features of anaphase chromosomes are more Rabl-like (Fig. 4d). Intriguingly, during anaphase, regions surrounding centromeres of chromosomes appeared to be more elongated as compared to interphase^G1^. Perhaps, the axial elongation of pericentromeric regions facilitates clustering of centromeres of different chromosomes in a confined nuclear space. To quantify these changes, we estimated the radius of gyration (R_g_) and asphericity (Fig. 4e) of polymer models of individual chromosomes at both stages. Interphase^G1^ chromosomes exhibited lower R_g_ and asphericity values, indicating more globular properties at interphase^G1^ than at anaphase. Thus, the polymer modeling helped us conclude that the shift from the non-Rabl configuration to the Rabl-type is associated with large-scale restructuring of chromosome organization.

**Fig. 4:**
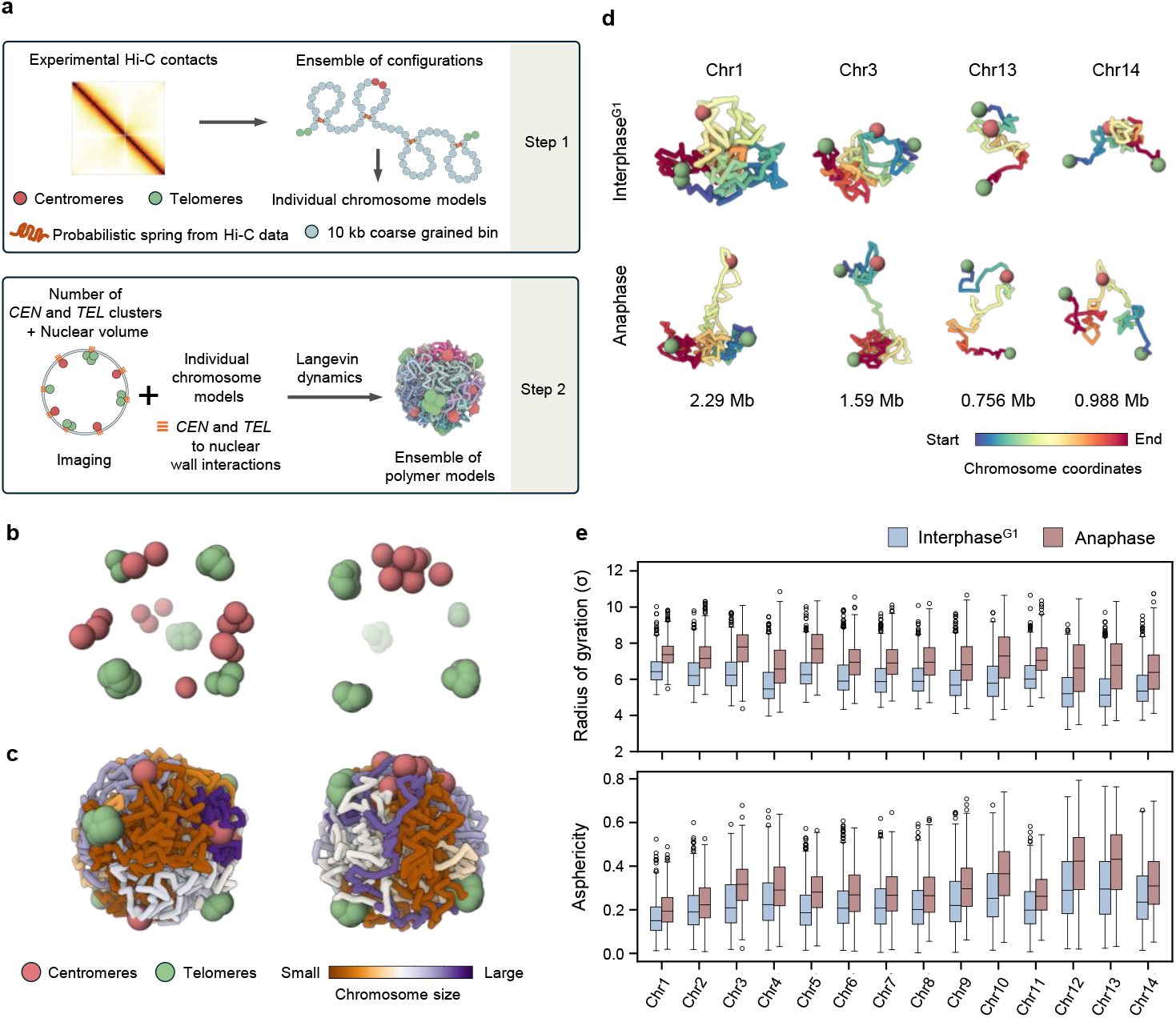
Polymer modeling of the *C. neoformans* genome. **a**, Schematic of the steps followed for generating a polymer model of the *C. neoformans* genome. **b**, Representative snapshot of simulations of interphase^G1^ and anaphase, highlighting only the spatial distributions of centromeres and telomeres. **c**, Snapshot of interphase^G1^ and anaphase whole genome polymer models. The individual chromosomes are colored according to increasing chromosome sizes. **d**, Snapshots of the polymer models of large (chr1; metacentric and chr3; acrocentric) and small (chr13; acrocentric and chr14; metacentric) chromosomes from interphase^G1^ and anaphase stages, colored from one end to the other (start to end coordinates). **e**, *Top*, Box plots of radius of gyration (R_g_) values of chromosomes at interphase^G1^ and anaphase. The higher the value, the more elongated the structure. *Below*, box plots of asphericity values estimated from the polymer models of chromosomes at the indicated stages. The lesser the value, the more globular the structures. The horizontal line inside the box represents the median, and the box denotes the interquartile range, 25% to 75%.

### Late-replicating centromere and telomere regions are positioned at the nuclear periphery

The spatial proximity of centromeres and telomeres and their compartmentalization from the rest of the genome during interphase^G1^ prompted us to test whether the genome is organized according to DNA replication timing as well. Several studies suggest a link between the 3D genome architecture and DNA replication timing^29^. Various parts of a genome are replicated at different time intervals within the S-phase. To estimate replication timing, we used sort-seq^30,31^ to measure the DNA copy number changes in replicating (S) cells relative to the non-replicating cells (G1). While in G1, the DNA copy number remains uniform, the copy number of different parts of the genome differs at S phase depending on the time of replication. Using this approach, we generated replication timing profiles of all 14 chromosomes of *C. neoformans* (Fig. 5a). A z-score >0 (methods) indicates early replicating and <0 indicates late replicating genomic segments. Centromeres and their flanks showed a dip in z-score values compared to other genomic regions. Similarly, subtelomeric regions (50 kb from the ends^32^) also exhibited a dip. These results helped us conclude that both centromeres and telomeres are late replicating, unlike the ascomycete fungi such as *S. cerevisiae, S. pombe*, and *Candida albicans*^33-35^, where centromeres are one of the earliest replicating regions, and telomeres replicate late in S phase. These observations, although unexpected, establish that the replication timing of centromeres and telomeres in *C. neoformans* is similar to metazoans^36,37^ rather than the yeast species analyzed to date. Next, we investigated if the early and late replicating regions fall within two different genomic compartments that were identified earlier.

**Fig. 5:**
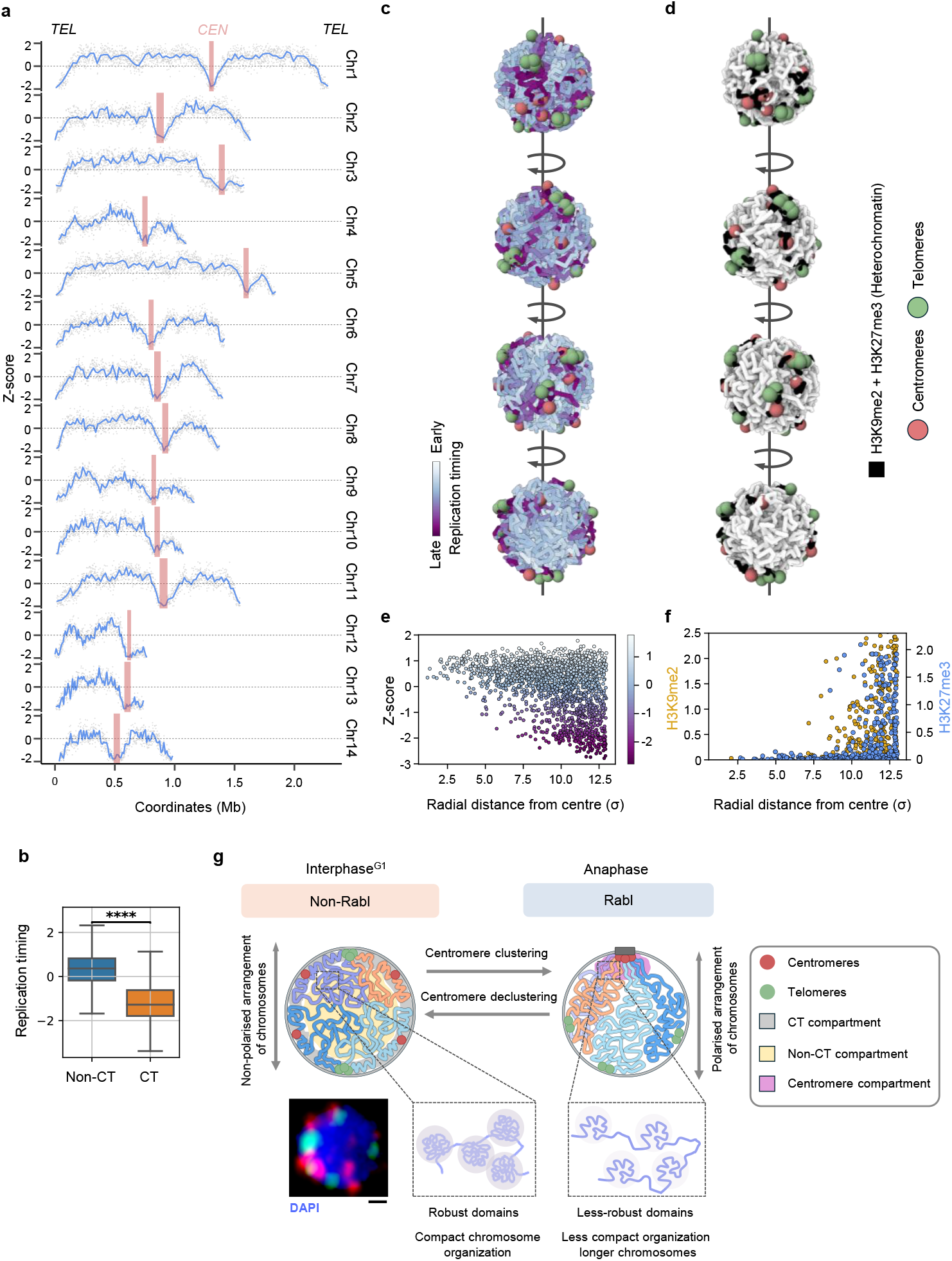
Late replicating heterochromatic regions are located at the nuclear periphery, insulated from the rest of the genome. **a**, DNA replication profile of all fourteen chromosomes. Red shaded boxes indicate centromeric regions. The x-axis represents chromosome coordinates in Mb, and the y-axis represents the z-score. Values >0 indicate early replication, and values <0 indicate late replication. Gray dots represent 1 kb bins. The blue line represents LOESS-smoothed z-score values. **b**, Box plot of z-score values (replication timing) of the indicated compartments. The Wilcoxon rank-sum test was used to check the significance. **c-d**, Representative snapshot of the polymer simulation of interphase^G1^ genome overlaid with, (**c**) the replication timing (z-score) values and, (**d**) the heterochromatin marks H3K9me2 and H3K27me3. Different views of the same structure are shown. The green and red spheres represent centromeres and telomeres, respectively. The blue-purple color gradient represents early to late replication timing. **e-f**, Scatter plot depicting the radial distance of the 10 kb beads of the polymer model from the center of the nucleus to the periphery with (**e**) replication timing and (**f**) ChIP enrichment profiles (log_2_ values) of H3K9me2 and H3K27me3 superimposed onto the beads. **g**, Model summarizing the changes associated with the reorganization of the genome during the shift between Rabl and non-Rabl states. In the model, a microscopy image of interphase^G1^ nucleus stained with DNA-binding dye DAPI and centromere and telomeres labelled with CENPA-mCherry and GFP-Pot1. Scale bar, 1 µm.

Indeed, the late replicating regions are found to be associated with CT compartments (Fig. 5b), and the early replicating regions belong to non-CT compartments. These observations provide experimental support that both physical and functional properties/states of chromatin dictate the interphase^G1^ genome organization. Next, we investigated if the observed late-replicating regions and heterochromatin are positioned towards the periphery of the nucleus. We mapped the replication timing (z-score) values and the ChIP enrichment values of genome-wide heterochromatin marks on the polymer model. This analysis revealed that most late-replicating and heterochromatic regions are positioned towards the periphery (Fig. 5c, d, e and f and Supplementary Videos 1 and 2). This was unlike what was reported in budding yeast, where a “replication wave” like structure^38^ is proposed to be present in which heterochromatic regions such as centromeres are enriched with early firing origins and telomeres are enriched with late firing origins that are arranged in a polarised fashion.

## Discussion

Here we report the 3D genome organization and DNA replication timing of the human pathogenic fungus *C. neoformans*. Using Hi-C, microscopy, and polymer modeling approaches, we provide evidence that the *C. neoformans* genome dynamically switches between two major chromosome configurations, the non-Rabl and Rabl, during the cell cycle. Our results demonstrate spatial positioning of centromeres and telomeres influences the overall organization of the chromosomes. Microscopic observations confirm that interphase^G1^ chromosomes in *C. neoformans* adopt the non-Rabl arrangement as centromeres and telomeres, occupy spatially proximal positions at the nuclear periphery (Fig. 5g). This is in striking contrast to the Rabl configuration observed in most ascomycete yeasts^13-15,17,39-42^. We show that centromeres dynamically cluster and decluster during the cell cycle, whereas telomeres remain in subclusters all throughout. The Hi-C data strongly support the previously reported microscopic observation^9^ that the centromeres gradually cluster in *C. neoformans* during the cell cycle progression. Interphase^G1^ Hi-C data indicate that pericentromeric and subtelomeric regions form separate CT compartments and avoid interactions with the rest of the genome. As a result, an interphase^G1^ chromosome is organized into three distinct blocks: a) euchromatin, b) the telomere block, and c) the centromere block. Centromere and telomere blocks constrain the overall organization of the interphase^G1^ chromosome by acting as an insulator. Though heterochromatin and euchromatin are well segregated in fungi^18,19^, the above mode of arrangement is unique in fungi studied so far. As expected, chromosomes undergo a dramatic reorganization as interphase^G1^ chromosomes transit from the non-Rabl to the Rabl-like configuration. In anaphase, only pericentromeric compartments exist, and the subtelomeric regions engage in long-range interactions with the arms of the chromosomes. Perhaps the polarised arrangement of centromeres and telomeres in the Rabl configuration masks the insulator effect of telomeres. The pericentromeric regions of anaphase are decondensed as compared to the rest of the genome. This is intriguing as in *S. pombe*, heterochromatin factors compact pericentromeric regions and promote inter-arm interactions between pericentromeric regions^18^. It will be interesting to investigate if the loss of pericentromere compaction promotes centromere clustering during anaphase in *C. neoformans*. Though there are differences in global chromosome organization, locally, chromatin in interphase^G1^ is folded into strong self-interacting domain-like structures reported in *S. cerevisiae, S. pombe*, and *V. dahlia*e^18,41,43^. The local chromatin organization also significantly differs between interphase^G1^ and anaphase in *C. neoformans*. As expected, the robust domain-like structures apparent in interphase^G1^ disappear in metaphase owing to compaction. On the other hand, these domain-like structures are reestablished during the transition from anaphase to interphase^G1^ of the next cell cycle.

A comparative study across several species of eukaryotes revealed that condensin II plays an important role in shaping the genome architecture across different species^1^. The presence of both condensin I and II promotes type-II/non-Rabl configuration, where chromosomes form territories. Most fungi lack condensin II subunits and thus favor the Rabl configuration^1^. Though *C. neoformans* lacks condensin II subunits, it can organize its chromosomes in a non-Rabl configuration. A recent study in *Arabidopsis thaliana*^44^ reported that condensin II and the linker of the nucleoskeleton and cytoskeleton (LINC) complex proteins help establish the non-Rabl configuration in interphase cells after exiting from mitosis. It has been shown in *C. neoformans*^9,45^ that both microtubules and a LINC complex protein, Sad1, are required for timely centromere clustering. Similarly, the cytoskeleton may regulate the clustering and declustering of centromeres, and to accommodate these dynamic changes, chromatin alters its folding properties independent of condensin II. Condensin II has been repeatedly gained or lost across many eukaryotic lineages^46^. Perhaps the presence of condensin II locks the genome in type II configuration, and its loss offers flexibility to change the genome organization to suit the needs of an organism.

In this study, we also provide evidence that, like other organisms, telomeres and subtelomeric regions are late replicating in *C. neoformans*. Whereas, unlike other yeasts studied, centromeres are late replicating. Late replicating centromere and telomere regions occupy spatially proximal positions (Fig. 5c) and are spatially sequestered as separate compartments from the relatively early replicating euchromatic regions, in contrast to the Rabl configuration of most budding yeasts. This raises the question of whether the relative position of centromeres with respect to that of telomeres within the nucleus influences centromere replication timing in general. For example, in *S. pombe*, chromosomes are arranged in the Rabl configuration, and centromeres bind to Swi6/HP1 and recruit DDK^47^, causing centromeres and their proximal regions to replicate early despite being heterochromatic. Meanwhile, at telomeres, the presence of PP1 phosphatase creates conditions that are refractory for the initiation of replication^48^. As a result, both centromeres and telomeres are present in different microenvironments that influence their replication timing^48,49^. Forced tethering of DDK to sub-telomeres did not advance the replication timing of telomeres like that of pericentromeric regions, suggesting that telomeric chromatin can hinder the recruitment of replication factors and delay its early replication^47,50^. Similarly, in *C. albicans*, a replication initiation protein Orc4 is enriched at the centromere cluster, and its concentration gradually reduces towards the telomeres. This proposed Orc4 gradient may, in principle, create conditions contributing to early and late firing of centromeres and telomeres associated origins respectively^51^. We hypothesize that telomeres may create an environment that promotes late replication of regions associated with them. In the Rabl configuration, centromeres are spatially separated from telomeres, whereas in non-Rabl, as observed in *C. neoformans*, the proximity of centromeres to telomeres may hinder early replication of centromeres. To verify if such a correlation exists in other organisms as well, we analyzed organisms in which the organization and replication timing of the genome have already been studied. In ascomycete yeasts possessing the Rabl configuration of chromosomes, centromeres replicate early, and telomeres replicate late. Intriguingly, in maize, with a Rabl-like arrangement, the centromeres replicate in mid S-phase^52^. On the other hand, chromosomes in humans, chicken DT40 cells, and *A. thaliana* adopt the non-Rabl arrangement, and centromeres replicate during mid to late S in these organisms^27,37,53-56^. These observations raise an intriguing link between the spatial positioning of telomeres and centromeres in the nucleus and replication timing of these heterochromatic domains across kingdoms of life. Altogether, our study highlights the conformational plasticity of the *C. neoformans* genome and maps the structural changes associated with the Rabl and non-Rabl conformational switching during the cell cycle. Most importantly, the conformation-switching behavior of *C. neoformans* chromosomes makes it a unique model system for studying the factors responsible for shaping the genome architecture in organisms that lack condensin II and sheds light on the evolutionary diversity of the 3D genome architecture.

## Acknowledgements

We thank members of the Sanyal laboratory for their support, discussion, and suggestions on the work. We thank R. Siddharthan, The Institute of Mathematical Sciences, Chennai, and L. Narlikar, IISER Pune, for the discussion and suggestions on the work. We thank V. Yadav, Duke University, for the discussion and comments on the manuscript. We thank K. Hardwick (University of Edinburgh) for the *MPS1* overexpression strain. We thank N. Nala and the in-house flow cytometry facility at JNCASR. We thank the NGS facility at NCBS and A. Pandit for sequencing Hi-C samples and for discussions on the DNA library preparation. This work was supported by the Anusandhan National Research Foundation (ANRF), Department of Science and Technology, Government of India (CRG/2023/001077) and intramural financial support from JNCASR to K.S. K.S. is a JC Bose National Fellow (JCB/2020/000021). S.D.P. acknowledges fellowship from ANRF (CRG/2023/001077), S.D. acknowledges fellowship support from the PMRF, Ministry of Education, India. R.P. acknowledges support from Sunita Sanghi Centre of Aging and Neurodegenerative Diseases (SCAN), IIT Bombay. The funders had no role in study design, data collection and analysis, decision to publish, or preparation of the manuscript.

