## Supplementary for "Organization principles of dynamic three-dimensional genome architecture associated with centromere clustering states"

This PDF includes:
Materials and Methods
Supplementary Figures 1-11
Supplementary Video legends 1 and 2
Supplementary Tables 1-3

### Materials and Methods

#### Strains and media growth conditions

A list of strains and plasmids used in this study is available in Supplementary Table 1. *C. neoformans* cultures were grown and maintained on YPD media (1% yeast extract, 2% peptone, and 2% dextrose) and YPG (1% yeast extract, 2% peptone, and 2% galactose) at 30°C unless otherwise specified. YPD+1M sorbitol plates were used for the transformation of *C. neoformans* cells. Approximately 100 µg/mL of nourseothricin (NAT, clonNAT, Werner BioAgents) was used for the selection of transformants. The primers used in this study to prepare constructs for generating and confirming the desired strains are listed in Supplementary Table 2.

#### Biolistic transformation

Biolistic transformation was performed as described previously[1]. Briefly, an overnight-grown 5 mL culture of *C. neoformans* was harvested and resuspended in 300 µL of sterile $dH_2O$. The cell suspension was spread on a YPD+1M sorbitol plate and allowed to dry. Gold micro-carrier beads (0.6 µm) coated with 2-5 µg of DNA, resuspended in 100% ethanol, were placed on a micro-carrier disk and allowed to dry. Dried disks containing DNA-coated beads were bombarded onto plates containing *C. neoformans* cells using the Biolistic® PDS-1000/He Particle Delivery System. Cells were then plated on YPD+NAT selection plates and incubated at 30°C for 3-4 days for transformants to appear.

#### Construction of *GFP-POT1* tagged strains

The fluorescent fusion protein of Pot1 was generated by cloning the 1.48 kb upstream homology region and promoter of *POT1* into the pVY7 plasmid containing the GFP NAT sequence. GFP was tagged to the N-terminal region of the *POT1* gene. The primers used to generate the clones are listed in Supplementary Table 2. The 1.48 kb homology region and 0.56 kb *POT1* promoter region were amplified from the *POT1* gene using H99 genomic DNA. Individual PCR fragments were purified using gel electrophoresis. The amplified regions were cloned using SpeI, SacI, and NcoI. The final clone was linearized using StuII and transformed into the CNVY101 background. The obtained transformants were confirmed by PCR.

#### Fluorescence microscopy

For widefield microscopy, a Zeiss Axio Observer 7 equipped with a 100x Plan Apochromat 1.4 NA objective, Colibri 7 LED light source, motorized XYZ stage, and PCO.edge 4.2 sCMOS camera was used. Images were acquired at 4 × 4 binning. Zen blue edition (2.3) software was used to control the microscope components and acquisition. For Airyscan imaging, a Zeiss LSM 880 Axio Observer confocal microscope equipped with a solid-state laser light source, 100x Plan Apochromat 1.4 NA objective, motorized XYZ stage, and Airyscan module was used to obtain images. Airyscan detection system was used to acquire images for centromere and telomere localization. Zen software (black edition) was used to control the microscope and process the Airyscan raw images. The number of centromere and telomere foci in unbudded cells was estimated using Fiji[2].

#### Super-resolution (SR) imaging of centromere and telomere localization

For SR imaging, log-phase *C. neoformans* cells were fixed using 4% paraformaldehyde in 1x PBS (pH 7.4) and incubated at RT for 20 min. The fixed cells were washed twice with 1x PBS, and 2µL of the cell suspension was placed on a microscope cover glass and imaged. For SR imaging, a Zeiss Elyra 7 lattice SIM[2] microscope equipped with a solid-state laser light source, Plan-Apochromat 63x 1.4 NA objective, motorized XYZ stage, and PCO.edge 4.2 sCMOS camera were used. Z-stack images were acquired using the leap mode at intervals of 0.11 µm. The raw images obtained were processed using the Zen software (Black Edition).

**Elutriation**

To isolate unbudded interphase$^{G1}$ cells, elutriation was performed using an Avanti J-26 XP (Beckman) in combination with a JE-5.0 elutriator rotor and a 40 mL elutriation chamber. The method was modified from[3] to suit the morphology of *C. neoformans* yeast cells. Log-phase asynchronous cell cultures were used for elutriation. An overnight-grown culture was used to inoculate 400 mL of 2% YPD at 0.05 $OD_{600}$/mL and grown overnight at 180 RPM, to reach 2.0 $OD_{600}$/mL. The following day, this culture was used to inoculate 2x 1 L YPD media (0.3 $OD_{600}$/mL) and was grown until ~ 1.5 $OD_{600}$/mL. The cultures were pelleted, washed twice with 1x PBS, and resuspended in 50 mL of 1x PBS. Clumped cells were resolved by sonication. Approximately $6 \times 10^9$ cells were pumped into the elutriation system at a flow rate of 14 mL/min and a rotor speed of 2500 rpm. Once the cells were loaded, equilibrium was achieved for 30 min, during which there was no output of cells from the chamber. To start eluting the unbudded G1 cells, the flow rate was gradually increased by 1 mL/min until the output became turbid. 1 L fractions of increasing flow rates were collected. The fraction containing unbudded cells was confirmed by FACS and microscopy and processed for Hi-C experiments.

**Synchronization of cells at metaphase and anaphase**

To obtain synchronous populations of metaphase and anaphase cells, the cells were first synchronized using hypoxia, which resulted in unbudded G2 arrest, as mentioned previously[4]. For metaphase arrest, hypoxia-synchronized cells were released by shifting to extensive aeration in YPGalactose (YPG) media to induce overexpression of Mps1. Mps1 has been shown to cause irreversible metaphase arrest in *C. neoformans*[5]. For the anaphase stage, hypoxia-synchronized cells were first arrested using thiabendazole (10 µg/mL) for 2 h, washed, and released into fresh YPD media. Synchrony was assessed using microscopy and flow cytometry.

**Hi-C**

This protocol was adapted and modified from previously published literature[6-8]. For Hi-C library preparation, approximately 50 $OD_{600}$ cells from three stages were suspended in 2% YPD and fixed with formaldehyde (3% final concentration) for 20 min, then quenched with 125 mM glycine final concentration for another 20 min at room temperature. The fixed cells were washed and resuspended in 1x Tris buffer (50 mM Tris-HCl pH 7.5, and 150 mM NaCl) with protease inhibitor cocktail and lysed using a Bio-spec Bead Beater. The lysate was centrifuged, and the pellet was washed with 1x NEB DpnII buffer. The pellet was digested with DpnII, followed by 14 biotin-dCTP fill in (Jena Bioscience), ligation, decrosslinking, Phenol: Chloroform: Isoamyl alcohol purification, precipitation, and RNase treatment as mentioned in[6]. DNA was fragmented using a Covaris M220 focused ultrasonicator to produce fragments in the size range of 200–500 bp. Sheared DNA fragments were fractionated using AMpure XP beads and pulled down using MyOne Streptavidin C1 beads. On-bead DNA library preparation was performed using the NEBNext Ultra II DNA library preparation kit (E7645), as reported previously[6]. The generated libraries were sequenced using 150-bp paired-end reads on an

Illumina Novaseq 6000 machine (NGS facility, NCBS, Bengaluru, India). The Hi-C sequencing read statistics are listed in Supplementary Table 3.

### Processing Hi-C data

Hi-C reads were aligned to the *C. neoformans H99* reference genome (ASM1180120v1)[9] and processed using the HiC-Pro (3.1.0) pipeline[10]. Fastq reads were mapped by Bowtie2[11] (2.3.5.1) using default parameters. Multiple hits, duplicates, and singletons were removed. The read pairs were assigned to the ligated restriction site DpnII, and invalid pairs were removed. Each replicate was processed independently, and the valid pairs were merged as one file and used for downstream analysis (refer to Supplementary Table 3 for read-pairing statistics). The valid pairs generated were converted to cool format and used to create balanced mcool files of resolutions 2, 4, 8, 16, and 32 kb using cooler[12]. Visual inspection of contact maps revealed structural changes associated with the genome of the strain used to generate the Hi-C maps. These regions were masked using cooler --blacklist option while generating balanced mcool files. The analyses presented in this paper were performed using replicate merged and masked mcool files.

### Visualization of Hi-C contact maps

The Hi-C contact maps and observed/expected correlated maps were plotted with $\log_{10}$, $\log_{2}$, and linear scales using the plotMatrix function from HiContacts[13] in R Studio 4.4.2.

### Plotting *P(s)* curves

The *P(s)* curves were plotted using 2 kb binned Hi-C matrices. Using cooltools[14], *P(s)* for the chromosomal arms was estimated.

### Calculation of eigen compartments

Eigenvalue decomposition of Hi-C matrices was performed using cooltools[14] at an 8 kb resolution. To determine compartment identity, the Pearson correlation between eigenvalues and gene density (computed over 8 kb windows for each chromosome) was calculated. The sign of each eigenvalue was adjusted based on this correlation: if the correlation was positive, the original sign was retained; if it was negative, the sign was inverted.

### Aggregate maps of centromeres and telomeres

Aggregate maps of inter/intra-chromosomal contacts involving centromeres and telomeres were generated using the aggregate function of HiContacts[13]. For off-diagonal contacts (*trans* contacts), 200 kb regions surrounding centromeres and 200 kb regions from the ends of the chromosomes (telomeres) were used. Aggregate contact maps binned at 8 kb resolution were plotted. A total of 182 pair-wise combinations of centromere and telomere p-arm interactions were plotted as aggregate plots. For on-diagonal interactions (*cis*) of centromeres and telomeres, Hi-C matrices binned at 2 kb resolution were used, and the aggregates were plotted as observed/expected maps using the detrend function from HiContacts. Aggregate maps surrounding the centromeres and telomeres were generated using the aggregate function from HiContacts. 8 kb binned matrices were used for this analysis and plotted using plotMatrix.

### Insulation score and domain calling

Insulation scores were calculated using FAN-C[15] on Hi-C matrices binned at 2 kb resolution using a 50 kb window. hicFindTADs[16] from HicExplorer was used to call the domains using a

2 kb binned Hi-C matrix of interphase$^{G1}$ stage using the following settings: --delta = 0.001, --minDepth 30000, --maxDepth 80000. The domains estimated using HicExplorer were aggregated across the genome using the FAN-C aggregate function and plotted as a heatmap (2 kb resolution). Hi-C contact maps, along with insulation profiles and domains, were plotted using PyGenomeTracks[17].

**ChIP-seq data processing**

The histone marks H3K27me3, H3K9me2, and control WCE ChIP-seq data used in the study were downloaded from GEO: GSE61550[18]. The raw sequence reads were aligned to the reference genome H99 using bowtie2[11]. The aligned files were sorted, duplicates were removed, and then sorted using samtools[19]. The ChIP-seq bam files were normalized to the sequencing depth. The log2 enrichment over control ChIP-seq data was then calculated for both histone mark data to generate bedGraph files over a window of 50 and 1000 bp using deepTools[20].

**RNA-seq**

RNA-seq data of G1 stage cells (0 min time point, post-elutriation released cells) were obtained from GEO: GSE80474[21]. The raw reads were aligned using the STAR rna-seq[22] aligner to the reference genome H99, and the respective aligned and sorted bam files were used to compare the gene expression patterns in the genome compartments.

**Circos plots**

Circos plots were generated using the Circos tool[23]. Hi-C data were processed using cooltools and HiCExplorer for analysis. Highly enriched 3D chromatin interactions (log2 ratio observed to the expected >3) are plotted as links. The eigenvalues were calculated using cooltools and are shown as separate tracks in the plot. The chip enrichment signal of histone modifications, that is, H3K27me3 and H3K9me2 over the control sample, is shown as two different tracks in the figure. Replication timing data are shown as another track in the plot.

**Sort-seq**

The protocol was adapted from a previously published report[24] with the following modifications. For sorting G1 and S phase cells, an overnight grown culture of *H99* cells was inoculated at 0.2 $O.D_{600}$/mL and grown for 5 h until the $OD_{600}$ reached 1. Approximately 20 O.D cells were washed twice with sterile water, fixed using 70% ethanol at room temperature for 1 h, and subsequently incubated overnight at 4°C. The cells were pelleted and washed twice with NS buffer (10 mM Tris-HCl pH 7.5, 250 mM sucrose, 1 mM EDTA (pH 8.0), 1 mM $MgCl_2$, 0.1 mM $CaCl_2$, and 7 mM β-mercaptoethanol). The cells were resuspended in 10 mL of NS buffer, treated with RNase A, washed with 1x PBS and stained with propidium iodide (PI) (HiMedia, Mumbai, India). The suspension was then incubated overnight in the dark at 4 °C. Before sorting, the stained samples were sonicated and sorted using FACSAria III (BD Biosciences). Approximately 15 million G1 and 10 million S phase cells were collected. As a control, 5 million total G1, S, and G2 cells were collected. The sorted cells were analyzed for purity (Supplementary Fig. 10 and Supplementary Fig. 11). Genomic DNA was isolated from both G1 and S phase cells using the bead beating method, subjected to library preparation using an Illumina kit, and the libraries generated were sequenced using 150-bp paired-end reads on the Illumina Nova-seq 6000 machine at Clevergene Inc., Bengaluru, India.

The Fastq files of the G1 and S phase samples were mapped using the Repliscope pipeline (https://github.com/DNAReplicationLab/Repliscope) with a few modifications. Briefly, the Fastq files were aligned using bowtie2 in paired-end and local modes, and the reads were binned using a 1000 bp window. Reads mapped more than once were not removed from the analysis by setting the MapQ threshold to 3 to include centromeric reads for downstream analysis. As reported previously[24-26], the read depth adjusted S/G1 ratio was calculated using makeRatio function of Repliscope, and the data were normalized to zero mean and 1 standard deviation, resulting in z-scores where values above and below zero indicate early and late replicating regions, respectively. The profiles were plotted using R Studio 4.4.2.

**Polymer simulation of individual chromosomes and the whole genome**

Individual chromosomes are composed of $N_i$ beads, each of size 10 kb and diameter $\sigma$. Intra-chromosomal beads ($\mathrm{i}$ and $\mathrm{j}$) are constrained to be in contact with probability $P_{ij}$ as reported in[27]. This was achieved in two steps, where the first step involved contact formation among the prominent contacts, followed by contact formation among all beads. After generating ensembles of 1000 polymer structures for each of the 14 chromosomes, ensuring that they accurately reproduced the experimentally observed intrachromosomal contact patterns, randomly selected configurations for each chromosome were placed together within a spherical confinement that mimicked the nuclear environment. Based on microscopy imaging of fluorescently tagged H4, the nuclear volume was set to approximately $4.29\,\mu\mathrm{m}^3$, which corresponds to a nuclear radius of about $1\,\mu\mathrm{m}$ under the assumption of a spherical shape. The radius of the spherical confinement was set to $14\sigma$, where $\sigma \approx 71.4\ \mathrm{nm}$. This choice of $\sigma$ yielded a confinement radius that matched the experimental nuclear size. The resulting chromatin volume fraction inside the nucleus is,

$$\frac{1914 \cdot \frac{4}{3}\pi(\sigma/2)^3}{\frac{4}{3}\pi(14\sigma)^3} = 0.087,$$

where $1914$ is the total number of beads. This volume fraction was in good agreement with the typical chromatin density observed in the nucleus[28]. The nuclear peripheral localization behaviour of centromeres and telomeres was incorporated into the model, using an attractive Lennard-Jones (LJ) interaction between the nuclear wall and both centromeres and telomeres, defined as:

$$E_{LJ}(r) = \begin{cases} 4\epsilon\left[\left(\frac{\sigma}{r}\right)^{12} - \left(\frac{\sigma}{r}\right)^{6}\right], & r < 2.5\sigma \\ 0, & r \geq 2.5\sigma \end{cases}$$

Here, r is the distance between any pair of interacting beads. While simulating wall-centromere, wall-telomeres, centromere-centromere ($\mathrm{cen}-\mathrm{cen}$) and telomere-telomere interactions ($\mathrm{tel}-\mathrm{tel}$), we took $\epsilon = \epsilon_{\mathrm{WC}}$ $\epsilon = \epsilon_{\mathrm{WT}}$, $\epsilon = \epsilon_{\mathrm{CC}}$ and $\epsilon = \epsilon_{\mathrm{TT}}$ respectively. All other beads interact via purely repulsive excluded volume interactions, where $E_{LJ}$ acts only where $\mathrm{r} < 2^{(1/6)}\,\sigma$.

From Hi-C data, we estimated the strong and weak interactions associated with centromeres and telomeres, and we accordingly tuned $\epsilon$ parameters and obtained configurations that are consistent with microscopy observations. Once the parameters are finalized, the system is equilibrated using Langevin dynamics simulations implemented in LAMMPS[29]. Here, we solved the Langevin equation with the above-mentioned interactions and thermal fluctuations.

**Calculation of distal contact index (DCI)**

To compare contact maps across different stages of the cell cycle and quantitatively understand how contact patterns vary, the Distal Contact Index (DCI) was computed[30]. The (normalized) local contact signal $C_l(i)$ and distal contact signal $C_d(i)$ were defined for each chromatin bin $i$ as:

$$C_l(i) = \frac{1}{N_l(i)} \sum_{j=1}^{N} P_{ij}\, \Theta(s_d - s_{ij}),$$

$$C_d(i) = \frac{1}{N_d(i)} \sum_{j=1}^{N} P_{ij}\, \Theta(s_{ij} - s_d),$$

where $P_{ij}$ denotes the contact probability between chromatin segments in bins $i$ and $j$, and $s_{ij}$ represents their genomic separation. The terms $N_l(i)$ and $N_d(i)$ denote the number of possible local and distal contact pairs, respectively, for bin $i$. The genomic distance threshold $s_d$ defines the boundary between local and distal contacts. We set $s_d = 70$ kb, based on the maximum domain size observed (Extended Fig. data 6). Contacts beyond this threshold are considered distal or nonlocal. Using the above definitions, the DCI for bin $i$ was computed as:

$$\mathrm{DCI}(i) = \log\left(\frac{C_d(i)}{C_l(i)}\right).$$

DCI is not well-defined in genomic regions that are difficult to sequence and lack reliable contact signals, such as the centromeric regions. A higher DCI value for a bin $i$ indicates the presence of contacts beyond typical domain sizes, which may signify the formation of long-range chromatin interactions.

**Calculation of compaction profiles**

The polymer radius of gyration is a conventional measure of compaction; alternatively, compaction can be defined based on contact matrices. Intuitively, higher compaction corresponds to higher contact probability values among genomic regions.

The average compaction $\langle P(k) \rangle$ of the $k$-th bin is defined as the mean contact probability within a symmetric window centered at $k$, of width $w$,

$$\langle P(k) \rangle = \frac{1}{N_w} \sum_{i=k-w/2}^{k+w/2} \sum_{j=k-w/2, j\neq i}^{k+w/2} P_{ij}$$

where $N_w$ is the number of non-diagonal terms.

**Radius of gyration and asphericity**

Chromosome shape characteristics, such as the radius of gyration and asphericity, were quantified using the gyration tensor. By calculating the eigenvalues of the gyration tensor ($\lambda_1$, $\lambda_2$, and $\lambda_3$), we derived the following shape descriptors:

$$\text{Radius of gyration (Rg)} = \sqrt{\sum_i \lambda_i}$$

$$\text{Asphericity} = \frac{3}{2}\left(\frac{\sum_i (\lambda_i - \bar{\lambda})^2}{(\sum_i \lambda_i)^2}\right)$$

where $\bar{\lambda}$ is the average of all eigen-values. Asphericity quantifies the deviation of the shape from a perfect sphere; it is zero for a sphere and increases as the shape deviates from that of a sphere.

**Polymer model visualization**

The simulation configuration snapshot images were visualized using the mol* viewer[31].

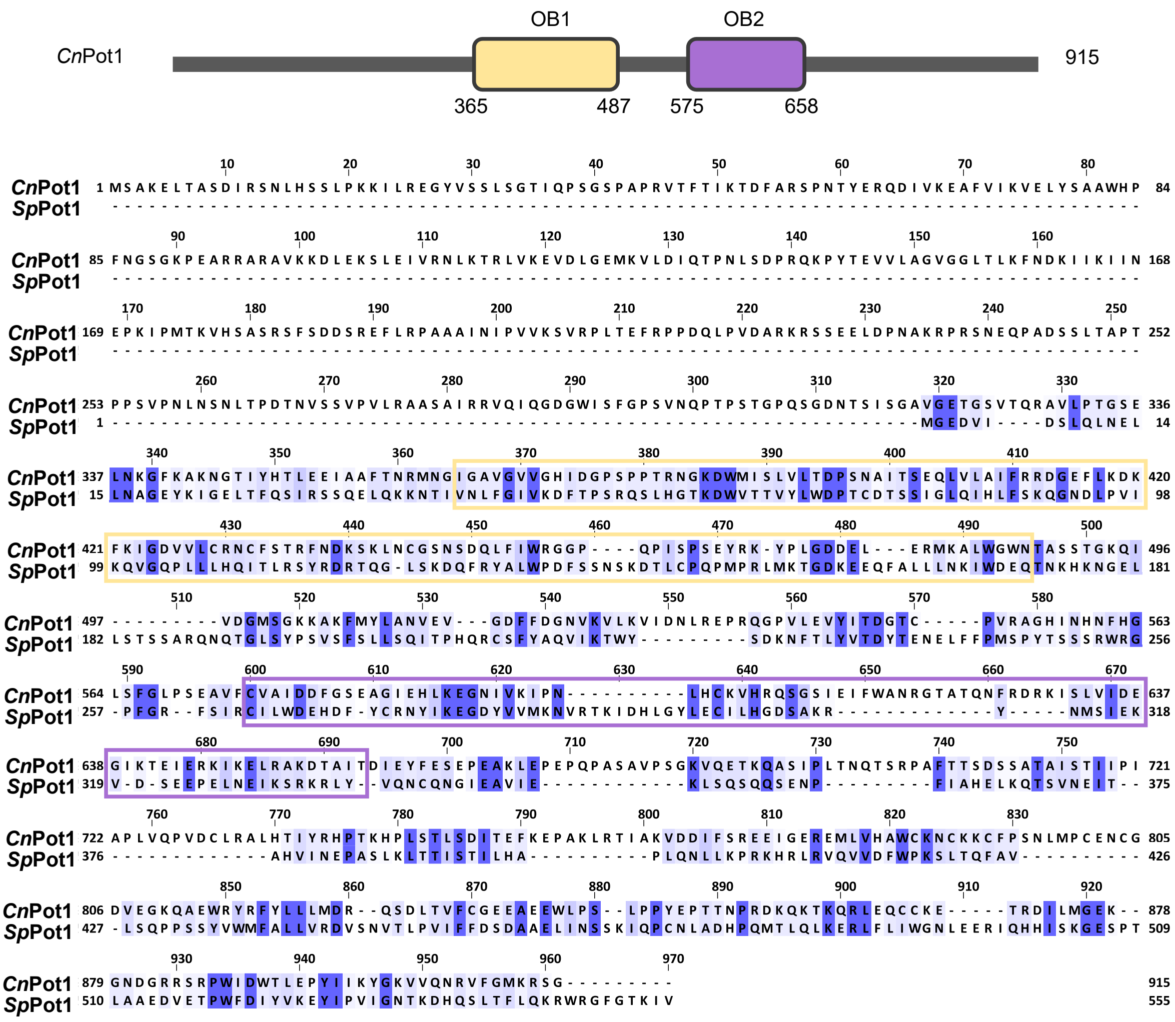


**Supplementary Fig. 1: Amino acid sequence conservation of *C. neoformans* Pot1 protein.** *Top*, Schematic representation of the domains present in *C. neoformans* Pot1 homolog. CnPot1 has two ssDNA binding domains (OB domains). *Below*, Pair-wise sequence alignment of *C. neoformans* Pot1 protein (CNAG_ 07661) and *S. pombe* Pot1 was generated using Clustal Omega[32] and annotated using Jalview 2[33]. An identity threshold of 30% was set as a cut-off for representing the sequence identity in the MSA. The two OB domains (E-values 2.3e-09 and 3.5e-06, respectively) corresponding to the CnPot1 are highlighted in yellow and purple in the MSA.

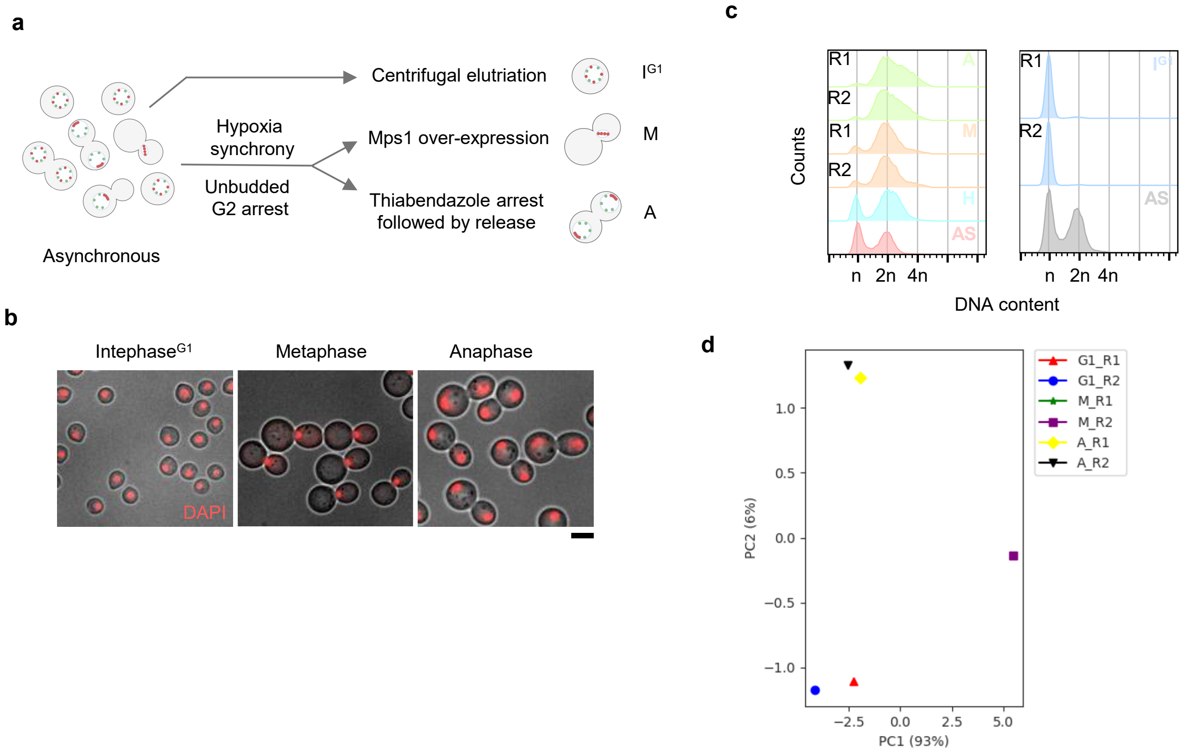


**Supplementary Fig. 2: Synchronization of *C. neoformans* cell cycle for obtaining populations of cells at different cell cycle stages. a**, Schematic of synchronization methods used for collecting cells at interphase ($I^{G1}$), metaphase (M), and anaphase (A) stages. **b**, Microscopy images of interphase$^{G1}$, metaphase, and anaphase cells labeled with DAPI to visualize DNA. Scale bar 10 μm. **c**, Flow cytometry profile to determine the stages of the cells obtained using synchronization. R1 and R2 represent replicates, H and AS represents hypoxia arrested cells and asynchronous control respectively. The x-axis represents cell counts, and the y-axis represents ploidy. **d**, Pearson correlation analysis of the Hi-C replicates of interphase ($I^{G1}$), metaphase (M), and anaphase (A) binned at 8 kb resolution using FAN-C pipeline.

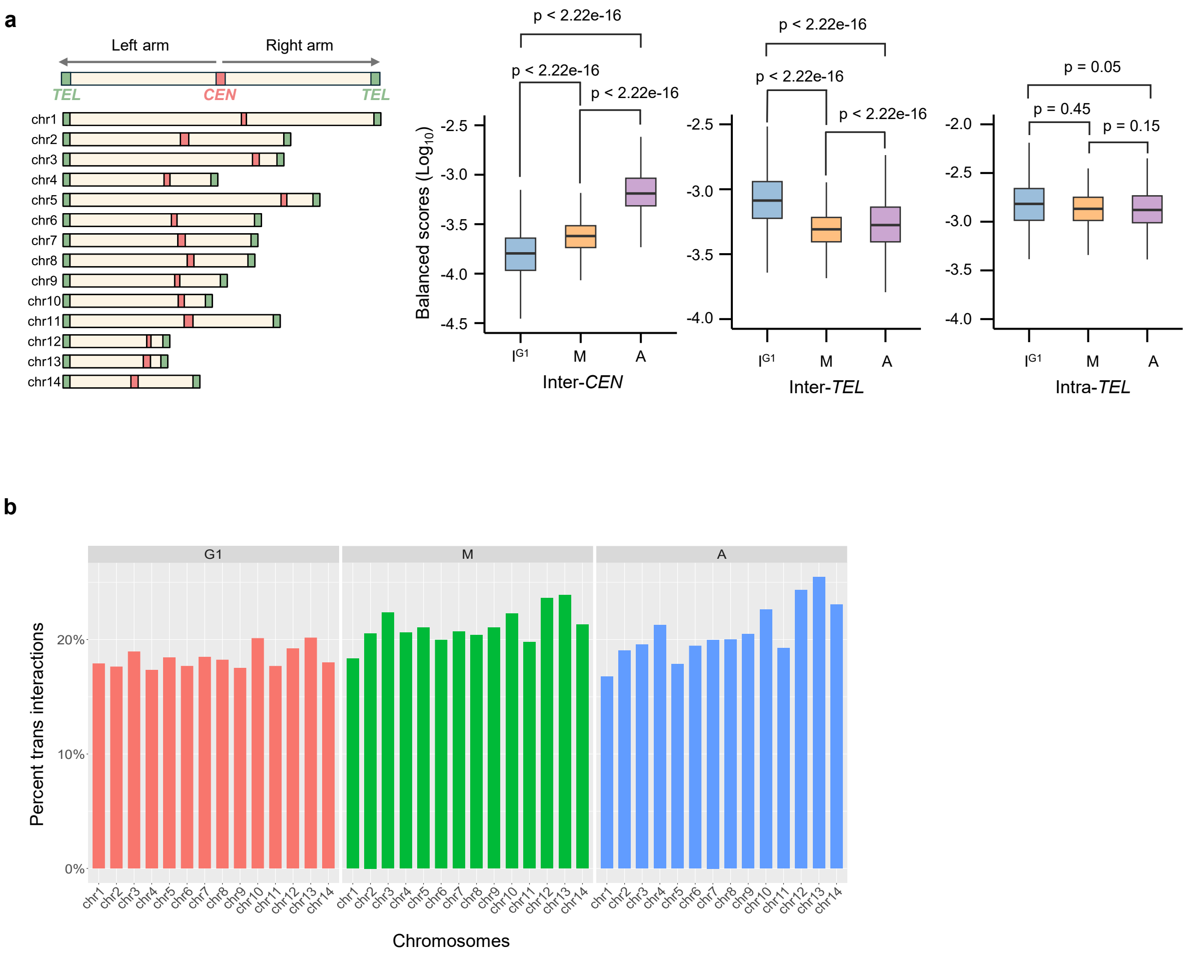


**Supplementary Fig. 3: Quantification of intra- and inter-chromosomal interactions. a**, *Left*, schematic of chromosomes with centromere, telomeres, and arms. *Right*, box plots representing quantification of the inter-centromeric and inter/intra-telomeric interactions among the chromosomes across interphase$^{G1}$, metaphase (M), and anaphase (A) stages. Centromeric regions with 50 kb pericentromeric regions, telomeres with 50 kb subtelomeric regions, were used to quantify the interactions. For inter-telomere interactions, regions corresponding to the left-arm of the chromosomes were used. The Wilcoxon rank sum test was used to test the significance. The horizontal line inside the box represents the median, and the box denotes the interquartile range, 25% to 75%. **b**, Percentage of *trans* (inter) chromosomal contacts across three different stages.

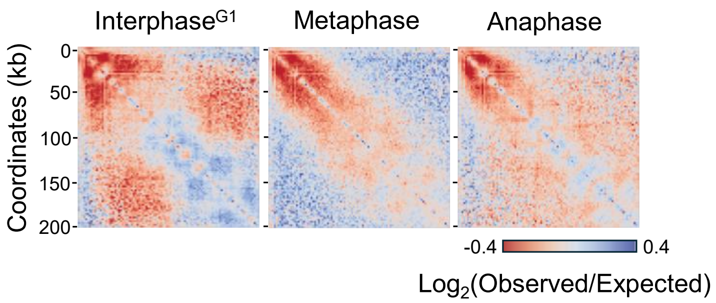


**Supplementary Fig. 4: Sub-telomeric regions are constitutively condensed.** Observed/expected aggregate Hi-C maps of *cis*-contacts of the subtelomeric region of the p-arm of all 14 chromosomes, binned at 2 kb. Color scale as indicated above. The color scale indicates enrichment of contacts (blue, least; red, maximum).

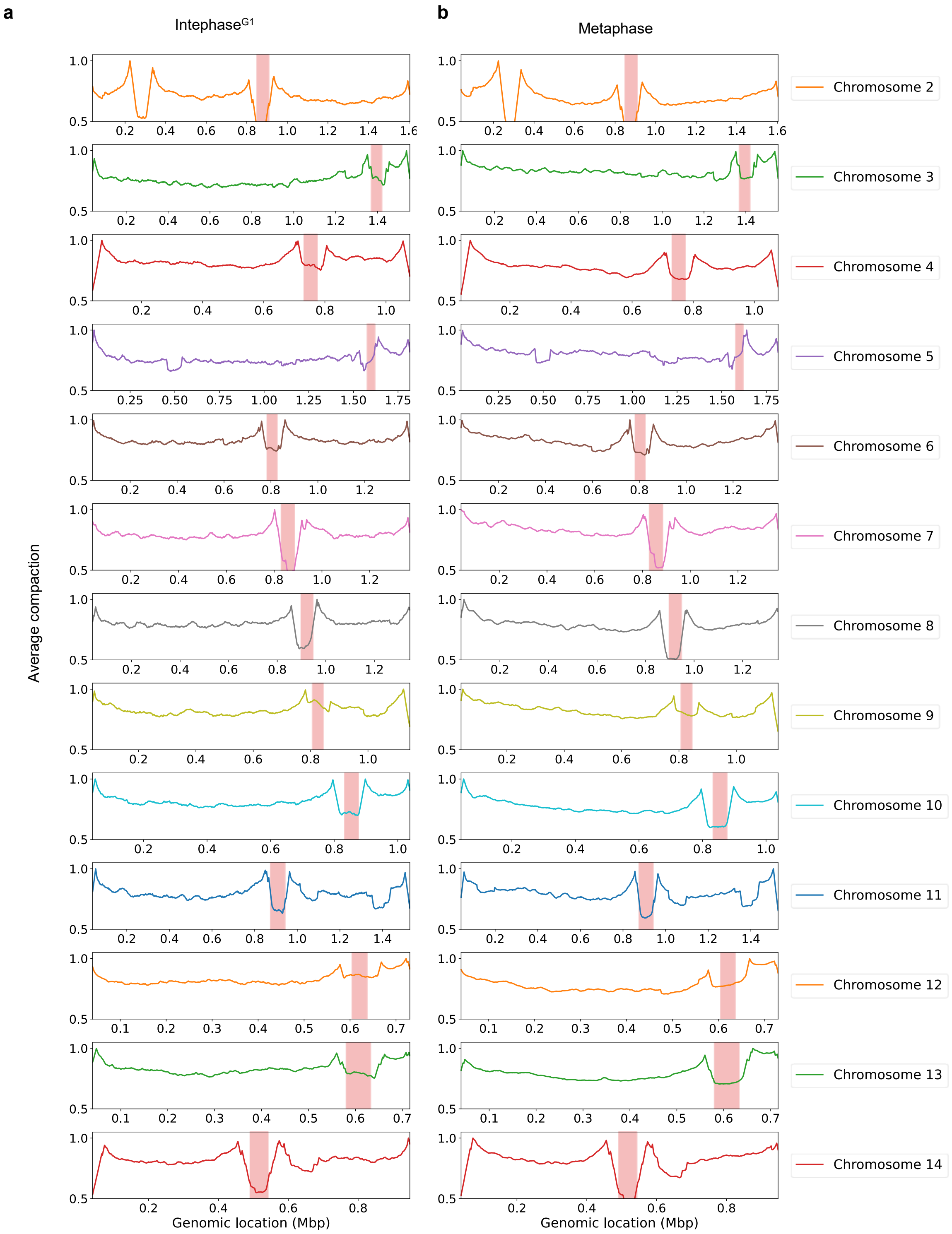
a
Intephase$^{G1}$
b
Metaphase
Average compaction
Genomic location (Mbp)
Genomic location (Mbp)
Chromosome 2
Chromosome 3
Chromosome 4
Chromosome 5
Chromosome 6
Chromosome 7
Chromosome 8
Chromosome 9
Chromosome 10
Chromosome 11
Chromosome 12
Chromosome 13
Chromosome 14

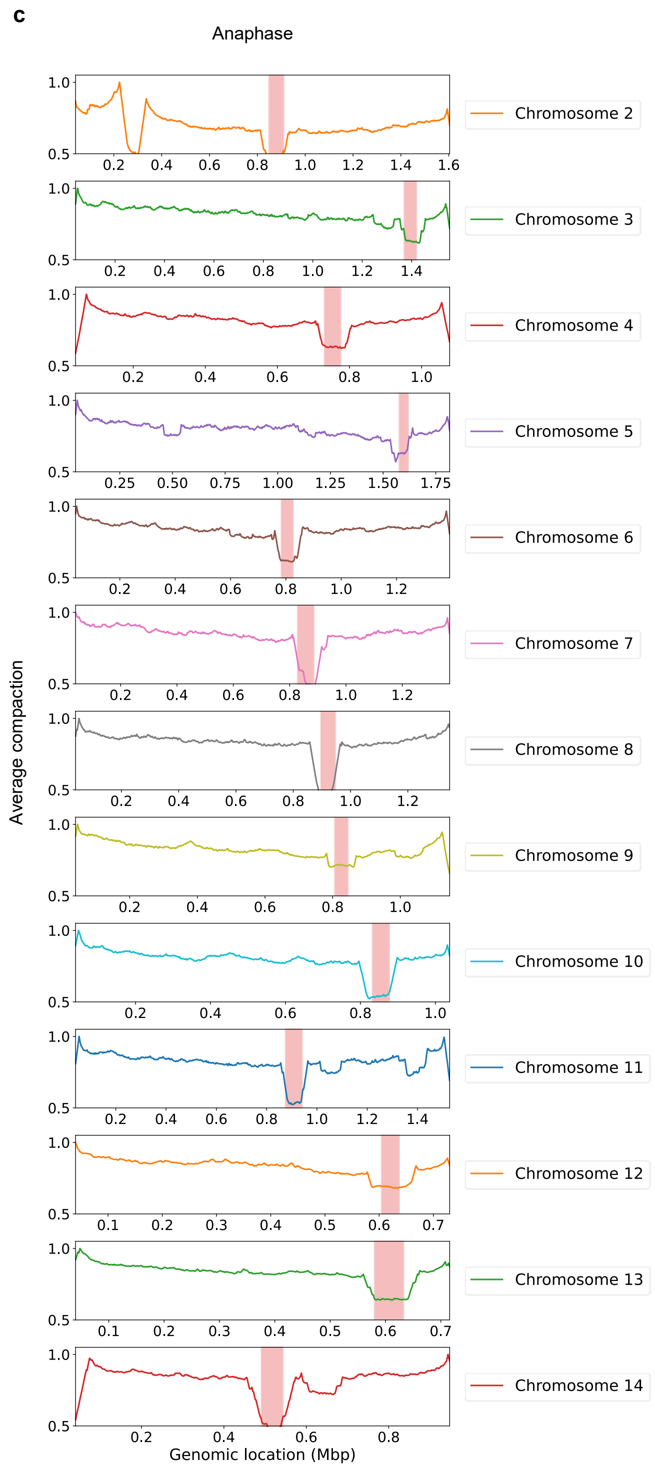


**Supplementary Fig. 5: Compaction curves. a-c,** Compaction profiles of chromosomes 2 to 14 for the indicated stages. The x-axis represents chromosome coordinates, and the y-axis represents average compaction. The centromere position of each chromosome is highlighted.

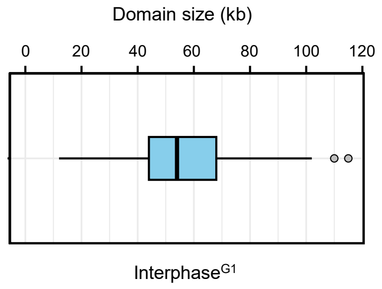


**Supplementary Fig. 6: Distribution of domain sizes at interphase$^{G1}$.** Box plot of domain sizes estimated from the domains called using hicFindTADs module of HiCExplorer. The vertical line inside the box represents median domain size, and the box denotes the interquartile range, 25% to 75%.

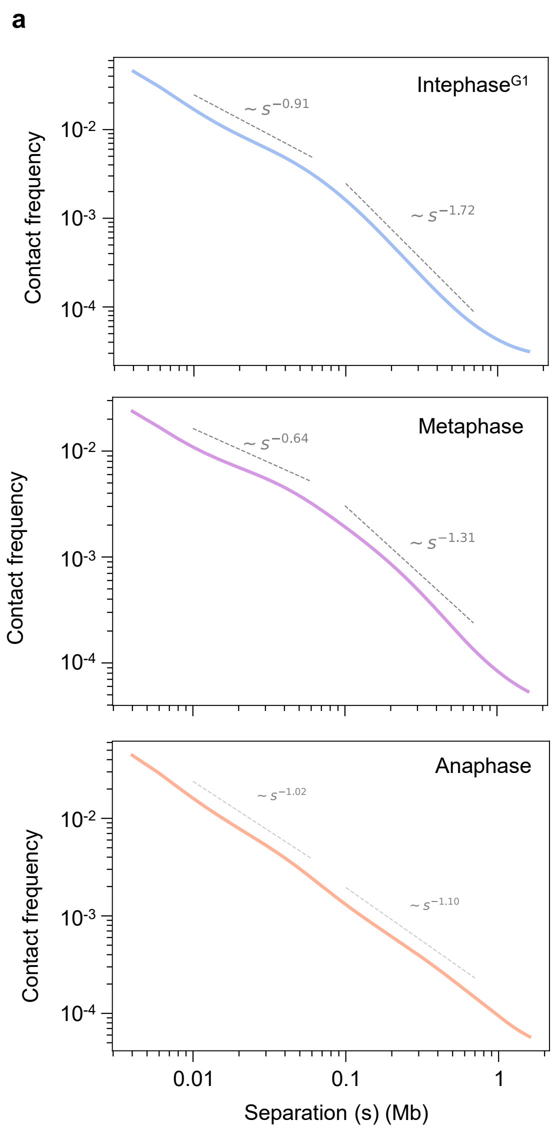
a
Intephase$^{G1}$
Metaphase
Anaphase
$\sim s^{-0.91}$
$\sim s^{-1.72}$
$\sim s^{-0.64}$
$\sim s^{-1.31}$
$\sim s^{-1.02}$
$\sim s^{-1.10}$
Contact frequency
$10^{-2}$
$10^{-3}$
$10^{-4}$
0.01
0.1
1
Separation (s) (Mb)

b

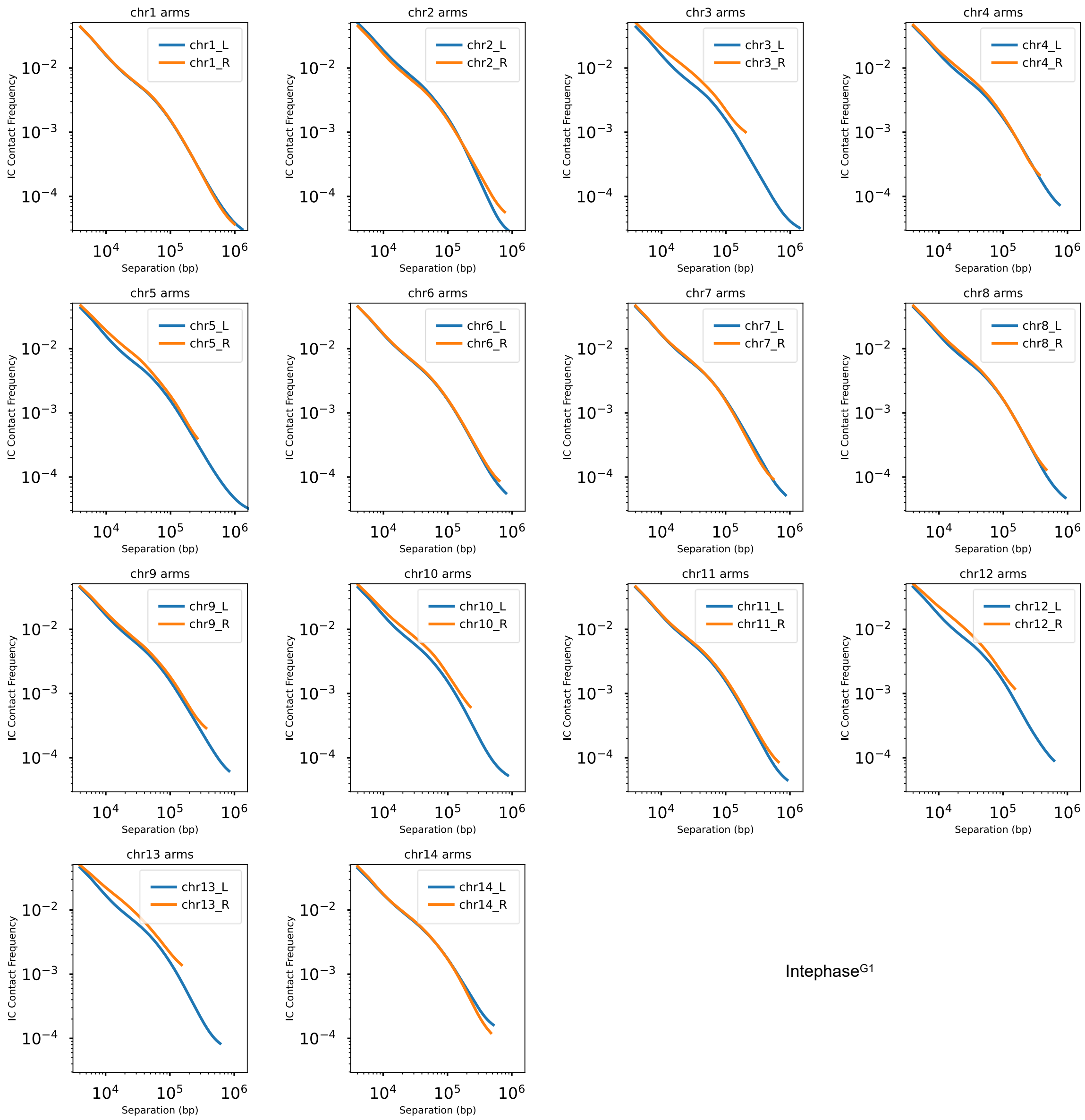
chr1 arms
chr1_L
chr1_R
chr2 arms
chr2_L
chr2_R
chr3 arms
chr3_L
chr3_R
chr4 arms
chr4_L
chr4_R
chr5 arms
chr5_L
chr5_R
chr6 arms
chr6_L
chr6_R
chr7 arms
chr7_L
chr7_R
chr8 arms
chr8_L
chr8_R
chr9 arms
chr9_L
chr9_R
chr10 arms
chr10_L
chr10_R
chr11 arms
chr11_L
chr11_R
chr12 arms
chr12_L
chr12_R
chr13 arms
chr13_L
chr13_R
chr14 arms
chr14_L
chr14_R
IC Contact Frequency
Separation (bp)
Intephase[G1]

c

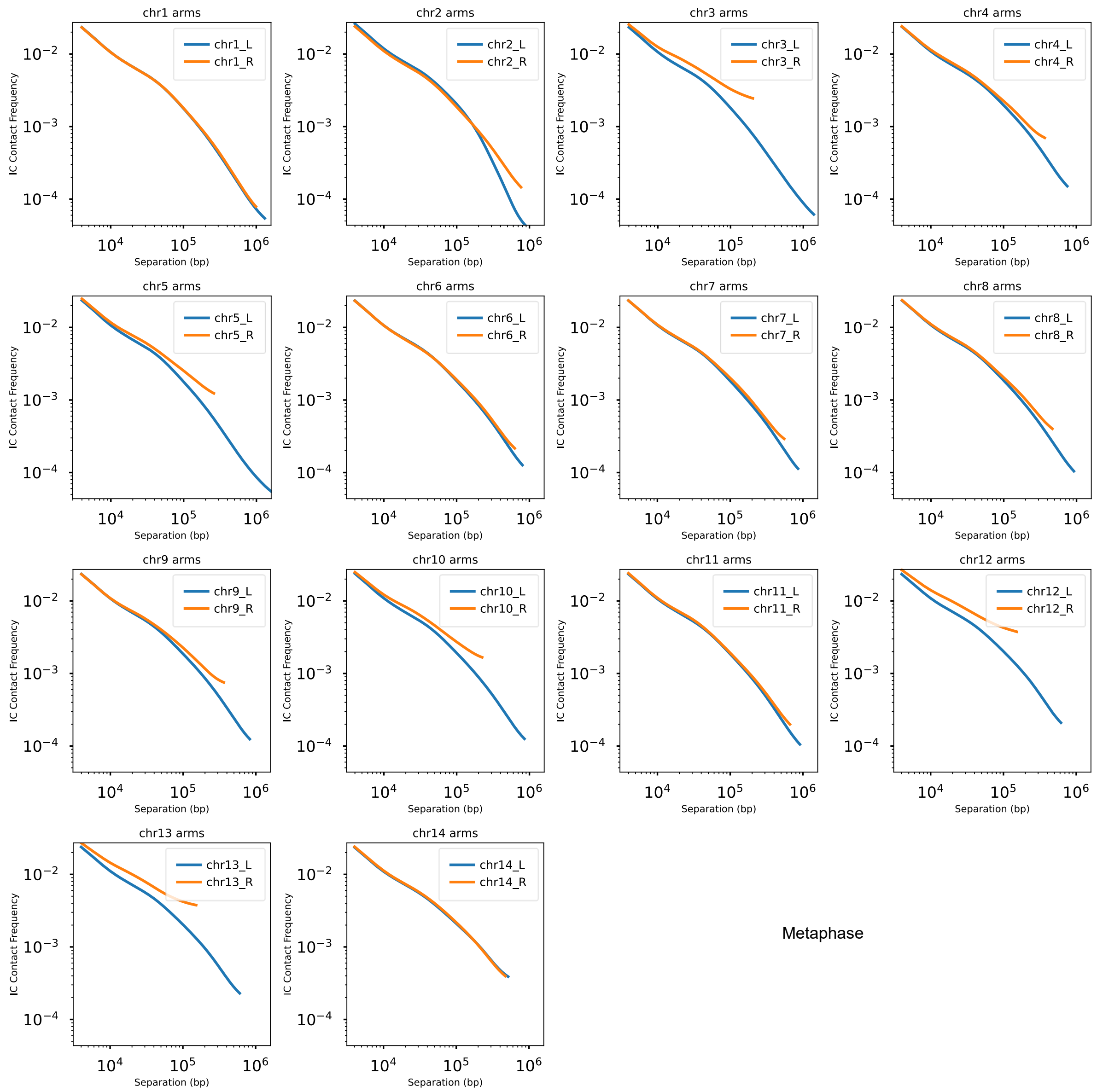
chr1 arms
chr1_L
chr1_R
chr2 arms
chr2_L
chr2_R
chr3 arms
chr3_L
chr3_R
chr4 arms
chr4_L
chr4_R
chr5 arms
chr5_L
chr5_R
chr6 arms
chr6_L
chr6_R
chr7 arms
chr7_L
chr7_R
chr8 arms
chr8_L
chr8_R
chr9 arms
chr9_L
chr9_R
chr10 arms
chr10_L
chr10_R
chr11 arms
chr11_L
chr11_R
chr12 arms
chr12_L
chr12_R
chr13 arms
chr13_L
chr13_R
chr14 arms
chr14_L
chr14_R
IC Contact Frequency
Separation (bp)
Metaphase

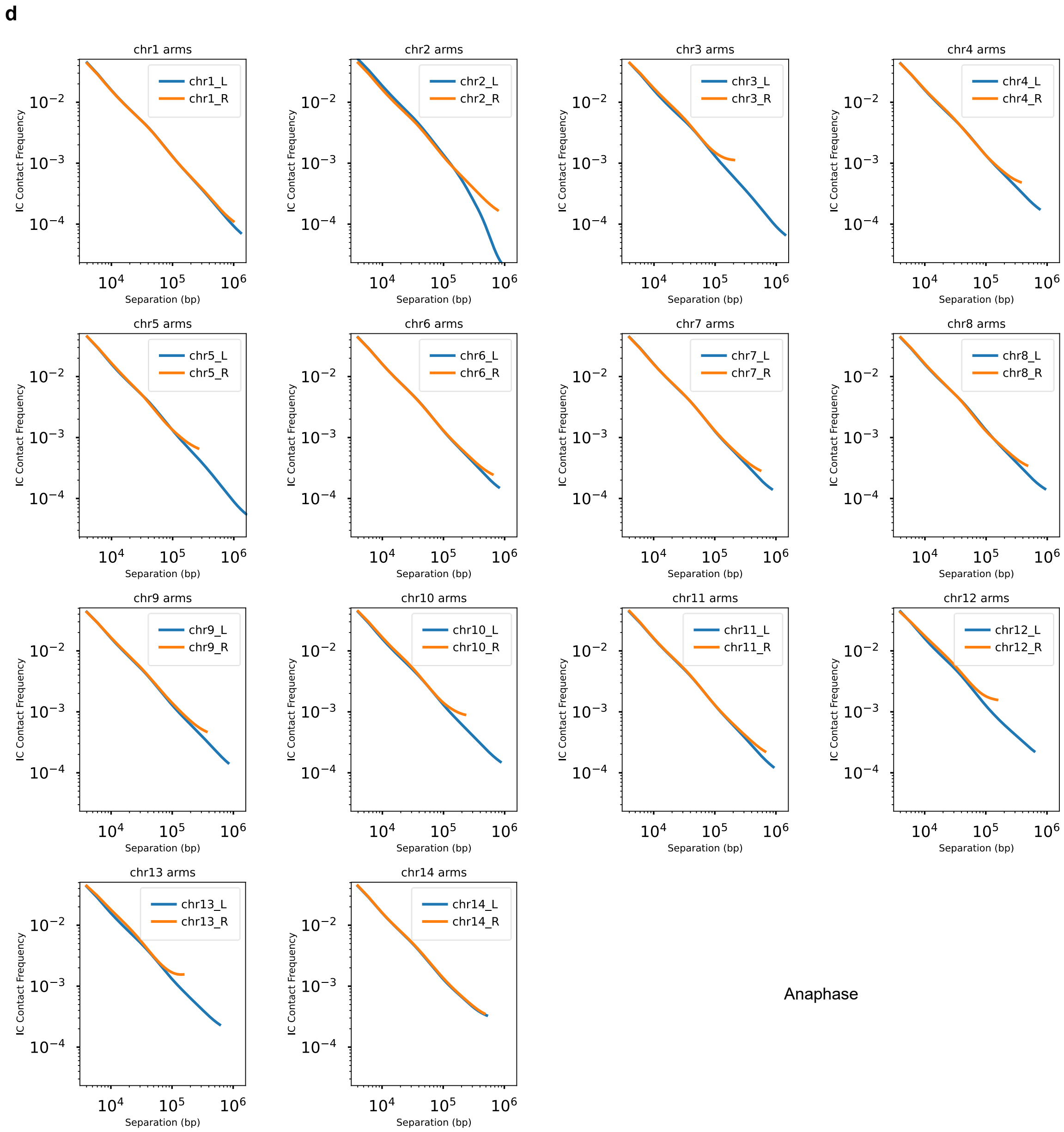


**Supplementary Fig. 7: Contact probability decay curves *P(s)* of individual chromosome arms at indicated stages. a,** Smoothened and aggregated contact probability decay curves of all the chromosomes of interphase[G1], metaphase, and anaphase. Dashed lines represent power-law scaling for the indicated regions. **b-d**, contact probability decay curves of individual arms of the chromosomes at the indicated stages. L and R represent left and right arms, respectively.

a b

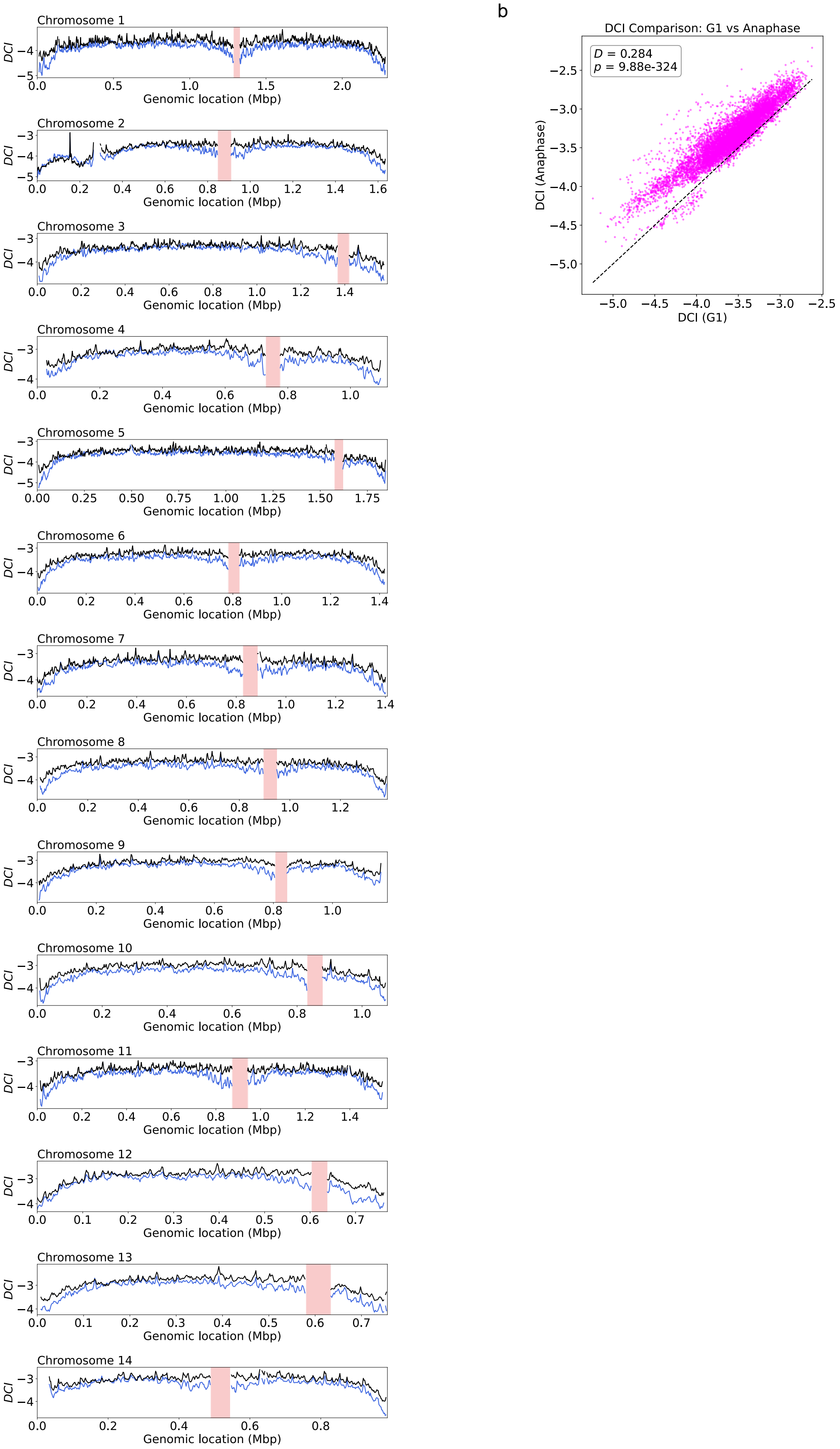

Chromosome 1
Chromosome 2
Chromosome 3
Chromosome 4
Chromosome 5
Chromosome 6
Chromosome 7
Chromosome 8
Chromosome 9
Chromosome 10
Chromosome 11
Chromosome 12
Chromosome 13
Chromosome 14
DCI
Genomic location (Mbp)
DCI Comparison: G1 vs Anaphase
D = 0.284
p = 9.88e-324
DCI (Anaphase)
DCI (G1)

**Supplementary Fig. 8: Distal chromatin index (DCI) of interphase$^{G1}$ vs anaphase chromosomes.** **a**, Distal chromatin index of the indicated chromosomes either at interphase$^{G1}$ (blue line) or at anaphase (black line) (see methods). Shaded regions represent centromeres, and these regions were omitted from the analysis. **b**, Scatter plot of DCI values of interphase$^{G1}$ and anaphase, highlighting an increase in distal contacts (above the diagonal) in anaphase. A two-sample Kolmogorov-Smirnov (KS) test indicates a statistically significant difference between the distributions (D = 0.284, $p \ll 10^{-4}$), suggesting that the DCI values are not identically distributed across the two cell cycle stages.

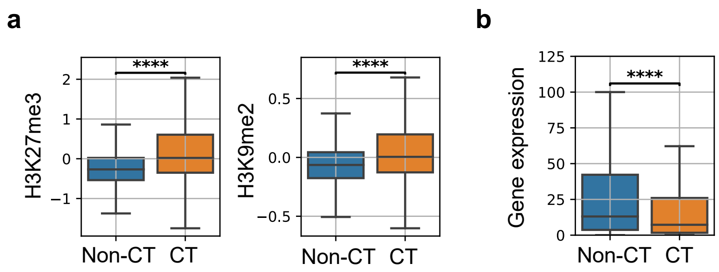


**Supplementary Fig. 9: Correlation of CT and non-CT compartments with heterochromatin marks, transcription levels, and replication timing. a,** Box plots representing $log_2$ fold change in the levels of H3K9me2 (left) and H3K27me3 (right) associated with the CT and non-CT compartments. **b**, Box plot highlighting changes in transcript levels of the indicated compartments. The Wilcoxon rank-sum test was used to check the significance.

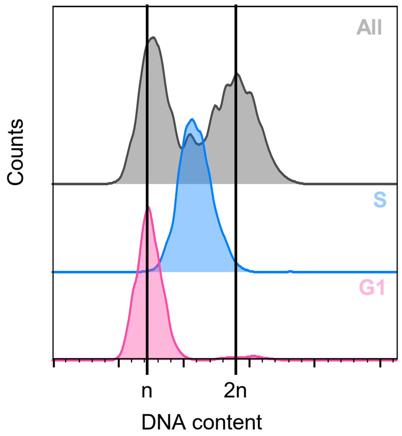


**Supplementary Fig. 10: Flow cytometry analysis of FACS-sorted G1 and S phase cells processed for the sort-seq assay.** Flow cytometry DNA content profile of the sorted cell populations stained with propidium iodide. ‘All’ represents the sorted cell population that includes G1, S, and G2/M stages of the cell cycle. x- and y-axes represent ploidy content and counts, respectively.

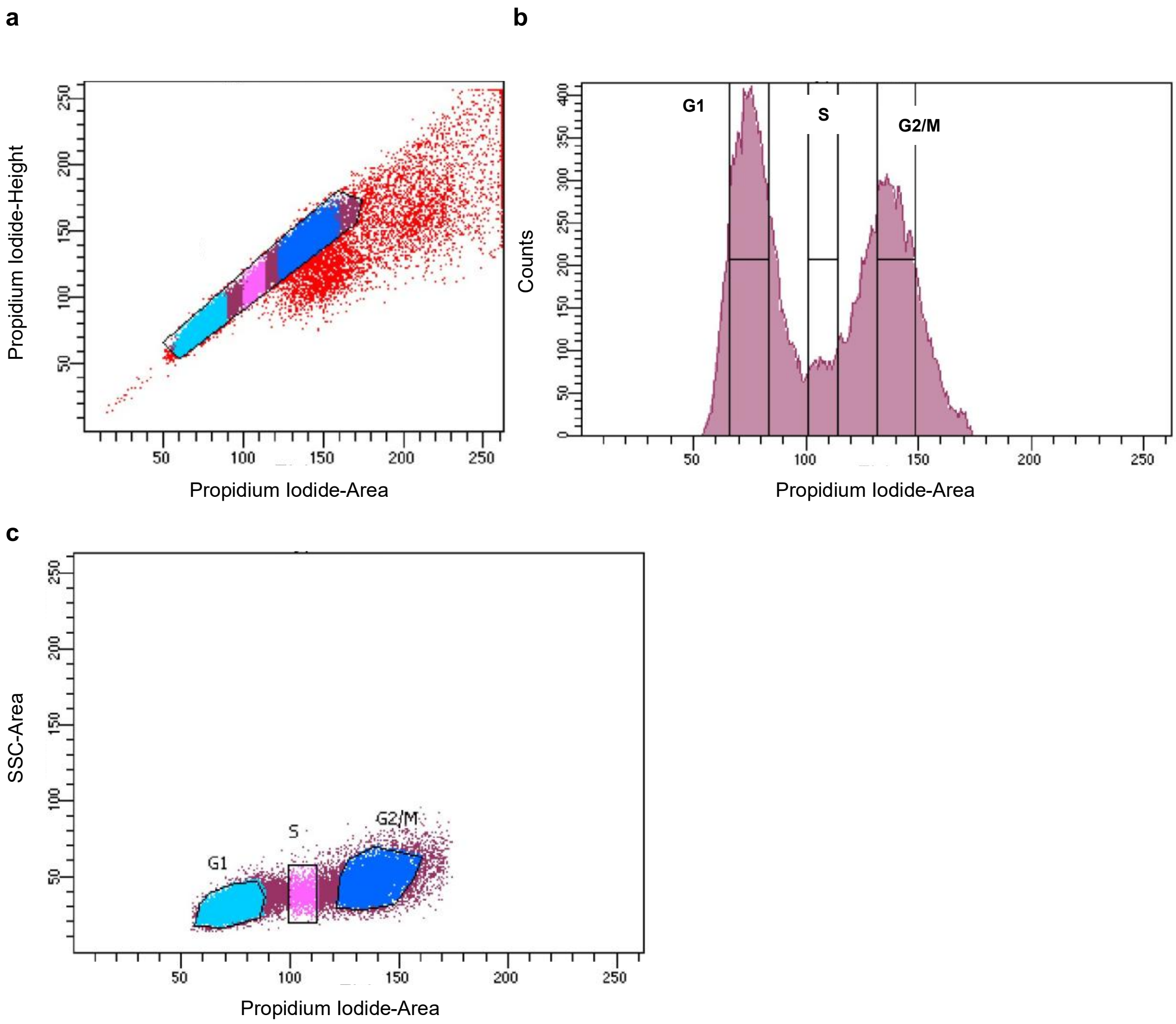


**Supplementary Fig. 11:** FACS plots showing (a) gating strategy to remove doublets using Propidium Iodide intensity and area and to include G1, S and G2/M the cell cycle stages (b) gating strategy to sort G1 and S phase cells using Propidium Iodide intensity (area versus height) (c) Distribution of the gated populations using side scatter area (SSC-Area) and Propidium Iodide area.

**Supplementary Video 1:** 3D model of interphase$^{G1}$ genome with DNA replication timing. Blue to purple gradient represents early to late replication timing. Centromeres and telomeres are represented as red and green spheres.

**Supplementary Video 2:** 3D model of interphase$^{G1}$ genome with heterochromatin marks. Regions containing H3K9me2 and H3K27me3 heterochromatic marks are colored black, and euchromatic regions are white. Centromeres and telomeres are represented as red and green spheres.

**Supplementary Table 1:** List of strains and plasmids used in this study.

| Name | Description | Source |
|---|---|---|
| H99 | *MATα* (wild-type) | [34] |
| CNSD230 | *MATα::POT1::POT1p-GFP-POT1-NAT (pSD18) mCherry-CENP-A-NEO (pLKB74)* | This study |
| IL08 | *MATα::GAL7p-myc-MPS1-NEO, chr14, SAFE HAVEN 7,* (pPEE36) *TUB1-GFP-NAT* | [5] |
| pLKB74 | *pXLI + CENP-Ap -mCherry-CENP-A (NEO)* | [35] |
| pSD18 | *H3p of pVY7 replaced with POT1p (SacI/NcoI) + POT1 homology region (SpeI)* | This study |

**Supplementary Table 2:** List of primers used in this study.

| | | |
|---|---|---|
| SDP87 | GGCACTAGTATGTCGGCCAAAGAGTTGAC | *POT1pr-GFP-POT1* N-terminal tagging |
| SDP88 | CGCACTAGTCGACTTATCGTTGAATCTGGTTG | |
| SDP89 | CTTGAGCTCGTGGAAGACATCCTCAGACAAC | |
| SDP90 | GTGCCATGGATTAGGCGATCGGACAGTTAAG | |
| SDP91 | CCACAAAGCCTTCATTCTCTC | *GFP-POT1* integration confirmation |

**Supplementary Table 3:** Hi-C read statistics generated using HiC-Pro pipeline.

| Sample | Total reads_R1 | Total reads_R2 | Mapped_R1 | Mapped_R2 | Valid reads (without duplicates) | Replicates combined |
|---|---|---|---|---|---|---|
| G1 Rep1 | 85200659 | 85200659 | 76911296 | 76355908 | 42256634 | 76249359 |
| G1 Rep2 | 76798649 | 76798649 | 68212998 | 67593142 | 34012207 | |
| M Rep1 | 61808992 | 61808992 | 55593017 | 55248597 | 32647599 | 71136612 |
| M Rep2 | 75151696 | 75151696 | 67033557 | 66556486 | 38493294 | |
| A Rep1 | 109006733 | 109006733 | 91893136 | 91099886 | 49054080 | 94583891 |
| A Rep2 | 97967058 | 97967058 | 82334004 | 81977035 | 45551531 | |
